# CXCL12 in late-stage osteoblasts and osteocytes is required for load-induced bone formation in mice

**DOI:** 10.1101/2022.08.25.505279

**Authors:** Pamela Cabahug-Zuckerman, Chao Liu, Pablo Atria, Cinyee Cai, Emily Fang, Shahar Qureshi, Rikki Rooklin, Cesar Ponce, Camila Morocho, Alesha B. Castillo

## Abstract

Increased physical loading of the skeleton activates new bone formation ensuring its ability to meet mechanical demands over time; however, the capacity of bone to respond to mechanical stimulation diminishes with age. Osteocytes, the cells embedded and dispersed throughout mineralized bone matrix, are master regulators of mechanoadaptation through recruitment of new bone-forming cells, the osteoblasts, via signaling to osteoprogenitors located on bone surfaces. We previously demonstrated that *in vivo* and *in vitro* mechanical stimulation significantly upregulated the chemokine C-X-C Motif Chemokine Ligand 12 (CXCL12) and its receptor, CXCR4, in osteocytes and bone lining cells, and that CXCR4 antagonism with AMD3100 attenuated *in vivo* load-induced bone formation. Here, we extended this work by showing that ablation of CXCL12+ cells and deletion of *cxcl12* in late-stage osteoblasts and osteocytes significantly attenuated *in vivo* load-induced bone formation in the mouse tibia. This bone loading phenotype was rescued by treatment with recombinant CXCL12. To address mechanism, we showed that *in vitro* deletion of *cxcl12* and *cxcr4*, separately, in bone marrow stromal cells resulted in significantly reduced osteogenic differentiation. Furthermore, CXCL12 treatment enhanced GSK-3b phosphorylation and β-catenin translocation to the nucleus, the former of which was partially blocked by AMD3100. Finally, CXCL12 synergized Wnt signaling leading to significantly increased total β-catenin protein and Axin2 expression, a Wnt signaling target gene. These findings together demonstrate that CXCL12 expression in late-stage osteoblasts and osteocytes is essential for load-induced bone formation, in part, by regulating osteogenic differentiation through activation of the Wnt signaling pathway.

**Significance:** Skeletal adaptation to mechanical loading is contingent on the recruitment of new osteoblasts to bone surfaces. CXCL12, a chemokine expressed by osteolineage cells, targets effector cells expressing its receptor CXCR4, including osteoprogenitors. Exogenous mechanical loading of mouse hind limbs upregulates CXCL12 in osteocytes, bone lining cells and marrow cells, while antagonizing CXCR4 led to significantly attenuated load-induced bone formation. Here, we show that CXCL12 expression in late-stage osteoblasts and osteocytes is required for load-induced bone formation. Treatment with recombinant CXCL12 rescued the bone loading phenotype suggesting that the CXCL12/CXCR4 signaling pathway may be a feasible drug target for promoting load-induced bone formation when exercise alone is insufficient to counteract low bone mass and osteoporosis.

## Introduction

Mechanical loading is a critical regulator of bone mass. Load-bearing exercise triggers new bone formation in a healthy skeleton (1, 2). However, skeletal mechano-responsiveness decreases with age leading to bone loss, osteoporosis and increased fracture risk. In addition, patients suffering from osteoporotic fractures often exhibit delayed bone healing and/or non-union (3). These age-related conditions converge on two critical deficiencies: a decrease in both number and osteogenic capacity of skeletal stem and progenitor cells (SSPCs) (4–7). To effectively treat this condition, we must better understand the cell and molecular mechanisms underlying load-driven osteogenesis, which appears to hinge on the terminally differentiated bone cells, the osteocytes, which act as bone mechanosensors (8–10). Osteocytes are individually embedded in small compartments throughout the bony matrix and their dendrites radiate into the matrix from the cell body, forming connections with neighboring osteocytes and an elaborate transport network through which they communicate with one another and with other cell populations located on bone surfaces (11–13). Given their ubiquitous nature, osteocytes have been recognized as master regulators of both bone formation and resorption in response to various stimuli including physical activity (14–16), disuse (17, 18), diet (19), reproductive status (20, 21) and injury (22, 23). Osteocytes respond to increased mechanical loading by releasing osteoprotegerin (24), which is an inhibitor of osteoclastogenesis, and pro-osteogenic factors such as insulin-like growth factor 1 (25), parathyroid hormone related protein (PTHrP) (26), polycystin 1 (27) and Wnts (28–30). They also down-regulate the expression of sclerostin (31, 32), an inhibitor of Wnt-signaling, an important regulatory pathway in osteogenic differentiation.

The CXC motif chemokine ligand 12 (CXCL12), first named stromal cell-derived factor-1 (SDF-1) (33) and shown to be essential during development (34), is an 8 kDa soluble protein abundantly expressed in bone marrow stromal cell populations (35, 36), and specifically by endothelial cells (37, 38), hematopoietic stem cells (38–40), osteoprogenitors (41, 42), adipogenic progenitors (41, 43) and osteoblasts (44, 45). CXCL12 is critical for stem cell recruitment (46, 47) and osteogenic differentiation (48–50) via coregulation with bone morphogenetic protein (48, 51) and Wnt signaling (42, 52). Our previous work showed that *in vivo* exogenous mechanical stimulation enhances expression of CXCL12 and its receptor, CXCR4, in osteocytes and bone lining cells, respectively, and that systemic antagonism of CXCR4 significantly attenuates load-induced bone formation in the mouse ulna (45). Similarly, others have shown that genetic deletion of CXCR4 in osterix (Osx)-expressing cells *in vivo* results in reduced skeletal size, lower bone volume and formation rates, disorganization of the growth plate, and inhibition of BMP-induced cell proliferation and osteoblast differentiation (53). Still unclear, however, is whether and how CXCL12 expressed specifically by osteoblasts and osteocytes affects bone structure, formation, and adaptation to mechanical loading.

In this study, we investigated the role of CXCL12 expression on basal bone metabolism and load-induced bone formation by evaluating mice in which CXCL12+ cells were ablated using diphtheria toxin A (DTA) and in separate mouse lines in which *cxcl12* was deleted from osteoblasts and osteocytes. Given the known role of Wnts in osteogenesis, we investigated crosstalk between the CXCL12/CXCR4 and canonical Wnt signaling pathways to clarify mechanism. Our results suggest a model in which mechanical loading up-regulates CXCL12 expression in osteoblasts and osteocytes, which then binds to and activates the membrane-bound CXCR4 receptor in osteoprogenitors, leading to osteogenic differentiation via synergism with the Wnt signaling pathway.

## RESULTS

### *Cxcl12* deletion results in lower trabecular bone volume and changes in some whole bone mechanical properties

To better understand the role of osteocyte- and osteoblast-specific CXCL12 expression on bone geometry and mechanical properties, tibias and femurs from adult 16-week-old mice harboring a CXCL12 gene deletion driven by the Col1α1 promoter (*Col1α1-cre;Cxcl12^fl/fl^*), the DMP-1 promoter (*DMP-1-cre;Cxcl12^fl/fl^*), separately, and their littermate controls (*Cxcl12^fl/fl^*_(*col1*)_ and *Cxcl12^fl/fl^*_(*dmp*)_, respectively) were examined. Trabecular bone microarchitecture in both *Col1α1-cre;Cxcl12^fl/fl^* and *DMP-1-cre;Cxcl12^fl/fl^* female mice was altered compared to their respective controls (**Fig. 1A**). *Col1α1-cre;Cxcl12^fl/fl^* females exhibited significantly lower trabecular bone volume (BV/TV) (−31%) and trabecular thickness (Tb.Th) (−14%) (**Fig. 1B**), while *DMP-1-cre;Cxcl12^fl/fl^* female mice exhibited significantly lower BV/TV (−19%) and greater trabecular spacing (Tb.Sp) (+15%) (**Fig. 1B**). No significant differences in trabecular microarchitecture were observed in males. To better understand the cellular mechanisms underlying the observed reduction in trabecular bone volume in females, proximal tibial metaphyseal sections were evaluated for osteoblast and osteoclast activity. Histological analyses of TRAP- and alkaline phosphatase (ALP)-stained thin-sections from 16-week-old female mice showed that the total number of osteoclasts were not significantly different between control and experimental mice (**Fig. 2A,B**). However, osteoblast perimeter was 40% (p=0.010) higher in female *Col1α1-cre;Cxcl12^fl/fl^* mice compared to their respective controls, with no differences observed in male *Col1α1-cre;Cxcl12^fl/fl^* mice or in *DMP-1-cre;Cxcl12^fl/fl^* female and male mice (**Fig 2C,E**). Femur length was significantly greater in female *DMP-1-cre;Cxcl12^fl/fl^* mice compared to their respective controls. No other geometric properties were altered upon *cxcl12* deletion (**Fig. S1A,B**). Yield stress was 12% lower in *DMP-1-cre;Cxcl12^fl/fl^* females compared to their respective controls (**Table S1**) and failure load was 14% higher in *Col1α1-cre;Cxcl12^fl/fl^* males compared to their respective controls (**Table S2**). No significant differences were detected in any other mechanical properties.

**Figure 1.**
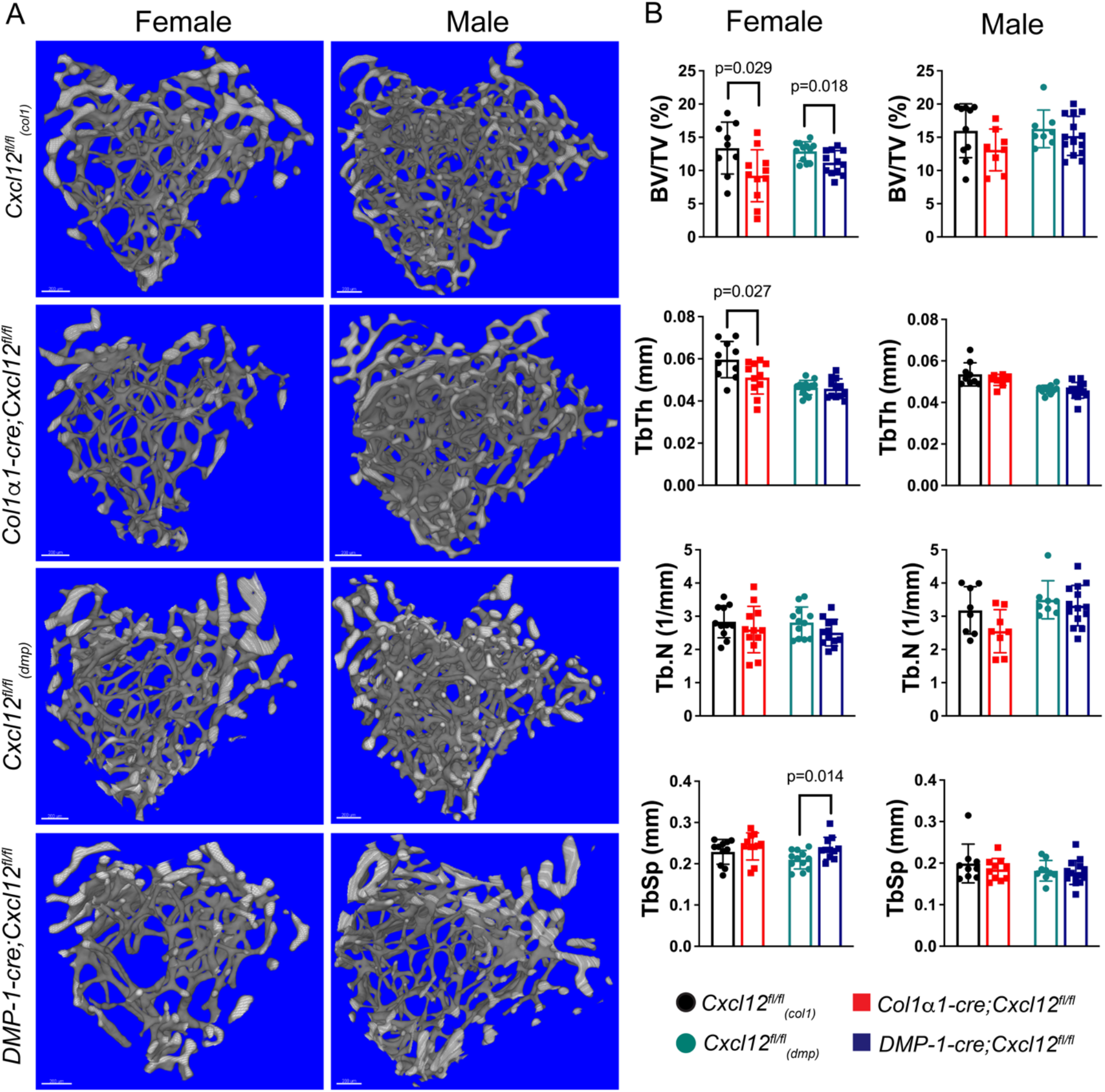
Trabecular bone volume and microarchitecture in *Col1α1-cre;Cxcl12^fl/fl^* and *DMP-1-cre;Cxcl12^fl/fl^* mice. (A) Representative μCT renderings of tibial metaphyseal trabecular bone in frontal view from 16-week-old *Col1α1-cre;Cxcl12^fl/fl^* and *DMP-1-cre;Cxcl12^fl/fl^* mice and their respective controls. Scale bar, 100 μm, n ≥ 7. (B) Quantification of trabecular bone volume fraction (BV/TV), thickness (Tb.Th), number (Tb.N), and separation (Tb.Sp). Data are presented as mean±SD.

**Figure 2.**
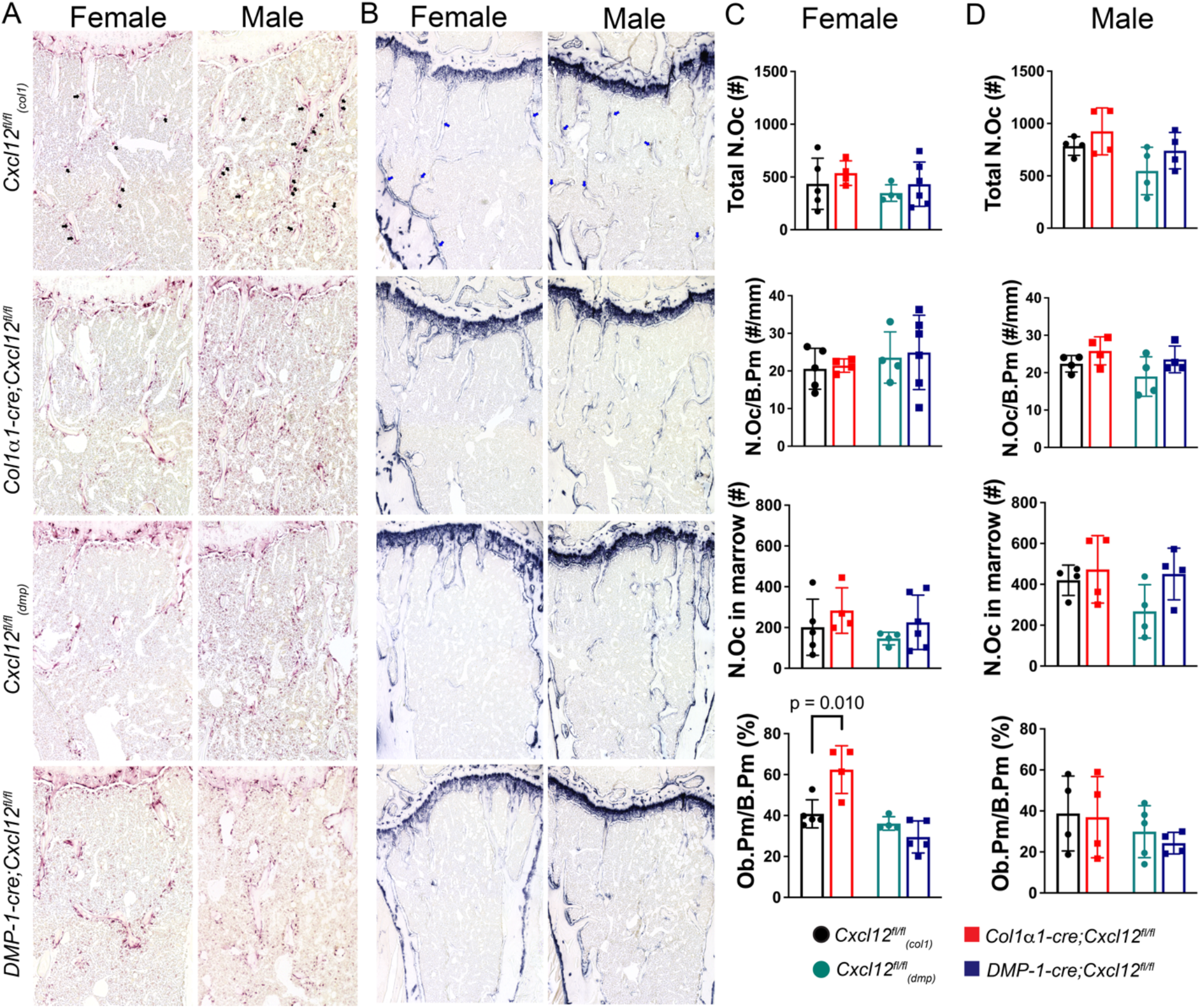
Osteoclasts and osteoblasts in trabecular bone in *Col1α1-cre;Cxcl12^fl/fl^* and *DMP-1-cre;Cxcl12^fl/fl^* mice. (A) Tartrate-resistant acid phosphatase (TRAP) staining in female and male 16-week-old tibias. (B) Alkaline phosphatase (ALP) staining in female and male 16-week-old tibias. (C) Osteoclast and osteoblast number and perimeter in female mice. (D) Osteoclast and osteoblast number and perimeter in male mice. N ≥ 4. Data are presented as mean±SD.

### CXCL12 signaling is essential for load-induced cortical bone formation

CXCL12 is highly expressed in SSPCs with expression decreasing as a function of differentiation into lineage specific phenotypes (36). We previously showed that CXCL12 mRNA expression was upregulated in cortical bone osteocytes and bone lining cells in the mouse ulna when subjected to anabolic exogenous loading (45). In the current work, the non-invasive murine axial compression tibial loading model (54) was used to study the role of CXCL12 in load-induced bone formation. First, CXCL12 protein expression was examined by thick-section confocal imaging of exogenously-loaded tibiae in CXCL12dsRed mice, which express DsRed-Express2 from the endogenous *cxcl12* mouse promoter. In tibiae experiencing normal ambulation (non-loaded), DsRed was visible primarily in marrow cells (**Fig. S2A, Video S1**). Tibiae subjected to anabolic exogenous loading exhibited DsRed expression in cells in the periosteum, cortical osteocytes and cells in the marrow (**Fig. S2B, Video S2**).

Next, static and dynamic bone formation parameters were calculated using *in vivo* fluorochrome bone labels in tibiae from *Col1α1-cre;Cxcl12^fl/fl^* and *DMP-1-cre;Cxcl12^fl/fl^* mice and their respective littermate controls during growth from 4 to 14 weeks old and in response to loading at 16 weeks old. No significant differences were detected in periosteal bone formation rates between ages 4 and 8 weeks (**Fig. S3 A,B,C)** and between ages 10 and 14 weeks (**Fig. S3 A, B, D**), respectively. To examine the role of CXCL12 in load-induced bone formation, a two-week load- and strain-matched loading protocol (**Fig. S4**) was applied to the right tibia of *Cxcl12-creER;Rosa26^DTA^, Col1α1-cre;Cxcl12^fl/fl^* and *DMP-1-cre;Cxcl12^fl/fl^* female and male mice and their littermate controls, with the contralateral tibia serving as an internal non-loaded control. Static and dynamic bone formation parameters were measured and relative (r) values (right minus left) were calculated. Relative mineralizing surface (rMS/BS) was significantly lower in *Col1α1-cre;Cxcl12^fl/fl^* and *DMP-1-cre;Cxcl12^fl/fl^* female mice compared to their controls (−75% and −57%, respectively) (**Fig. 3 A,B,C**). Relative mineral apposition rate (rMAR) was also significantly lower in DMP-1-cre;Cxcl12fl/fl female mice versus controls (−53%). These differences were reflected in significantly lower relative bone formation rates (rBFR/BS) in *Col1α1-cre;Cxcl12^fl/fl^* and *DMP-1-cre;Cxcl12^fl/fl^* female mice compared to their controls (−64% and −57%, respectively). Male *DMP-1-cre;Cxcl12^fl/fl^* mice exhibited a significantly lower rMS/BS (−70%) (**Fig. 3 A,B,C**), but not rMAR, resulting a significantly lower rBFR/BS (−64%). No differences were observed in male *Col1α1-cre;Cxcl12^fl/fl^* mice. These results are supported by data showing significantly lower rMS/BS (−71%) and rBFR/BS (−77%) in loaded tibiae from mice undergoing targeted ablation of CXCL12+ cells (**Fig. S5**) using selective expression of diphtheria toxin A (DTA).

**Figure 3.**
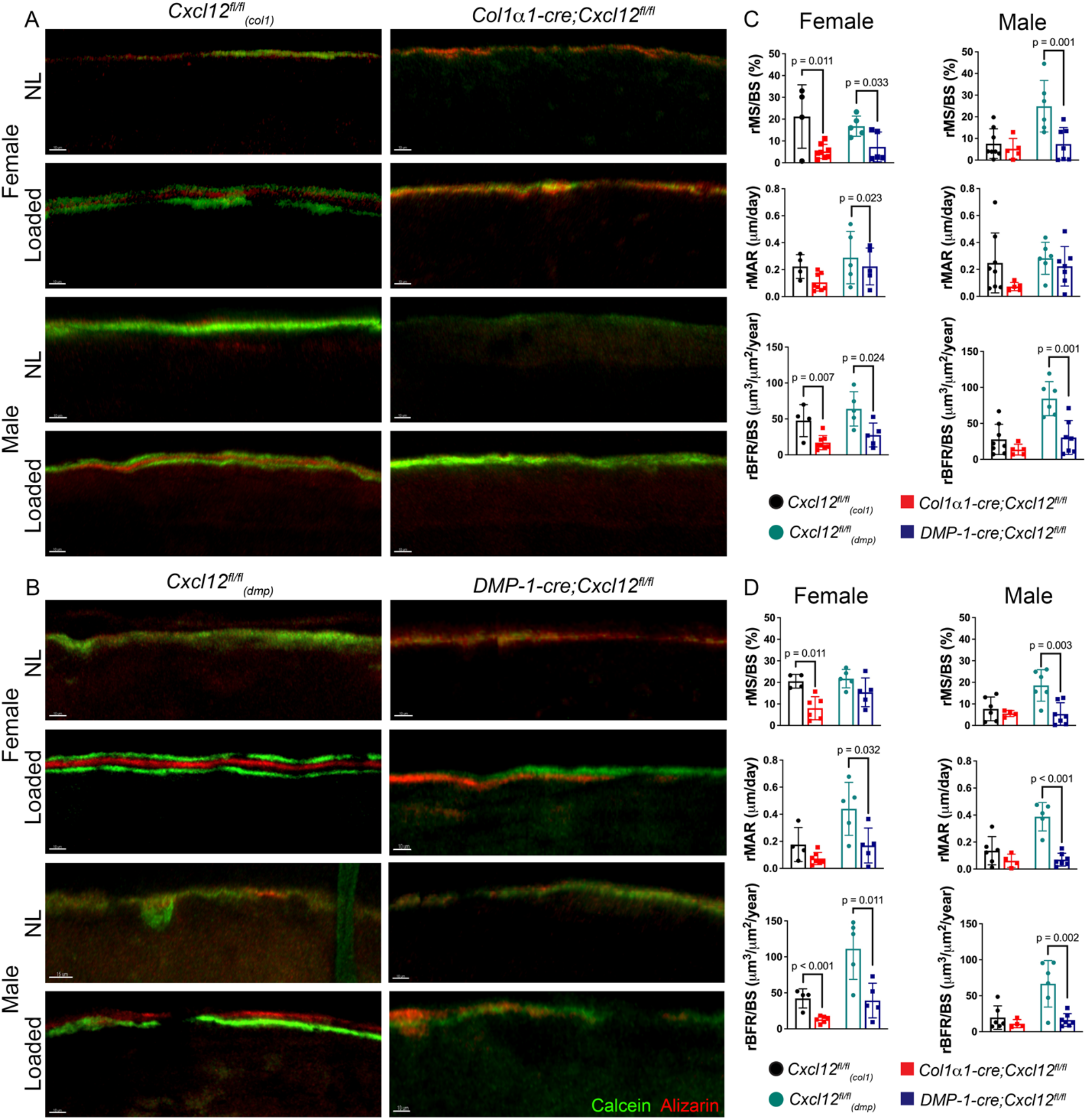
Load-induced bone formation in *Col1α1-cre;Cxcl12^fl/fl^* and *DMP-1-cre;Cxcl12^fl/fl^* mice. In vivo fluorochrome bone labels (calcein, green; alizarin, red) at the periosteal surface in loaded and non-loaded (NL) tibias from female and male (A) *Col1α1-cre;Cxcl12^fl/fl^* and (B) *DMP-1-cre;Cxcl12^fl/fl^* mice and their respective cre-negative littermate controls. Relative mineralizing surface (rMS/BS), mineral apposition rate (rMAR) and bone formation rate (rBFR/BS) between post-loading days (C) 5 to 12 and (D) 12 to 19. N ≥ 4. Data are presented as mean±SD.

### Exogenous CXCL12 treatment rescues the bone loading phenotype

To investigate whether CXCL12 treatment could rescue the bone loading phenotype of mice harboring a CXCL12 deletion, we administered recombinant human CXCL12 (R&D, #350-NS) encased in a biopolymer (BRTI, CellMate 3D # CM-1001) via subdermal injection. The injection was positioned adjacent to the bone surface at the mid-diaphysis of the right tibia in *DMP-1-cre;Cxcl12^fl/fl^* male mice on 1 and 8 days following the first day of loading (**Fig. 4A**). *DMP-1-cre;Cxcl12^fl/fl^* tibiae treated with CXCL12 and exposed to mechanical loading (CXCL12+L) exhibited full recovery of their osteogenic capacity in response to loading (**Fig. 4B**), as reflected in significant load-induced increases in MS/BS, MAR and BFR/BS relative to their internal contralateral non-loaded controls treated with vehicle alone (Veh+NL) (**Fig. 4C**). This result was similar to increases in absolute (**Fig. 4C, aqua bars**) and relative bone formation parameters (**Fig. 4D**) observed in *CXCL12^fl/fl^* control mice. No differences in bone formation parameters were detected between vehicle-treated and CXCL12-treated tibiae exposed to normal ambulation (non-loaded) (**Fig. 4E**).

**Figure 4.**
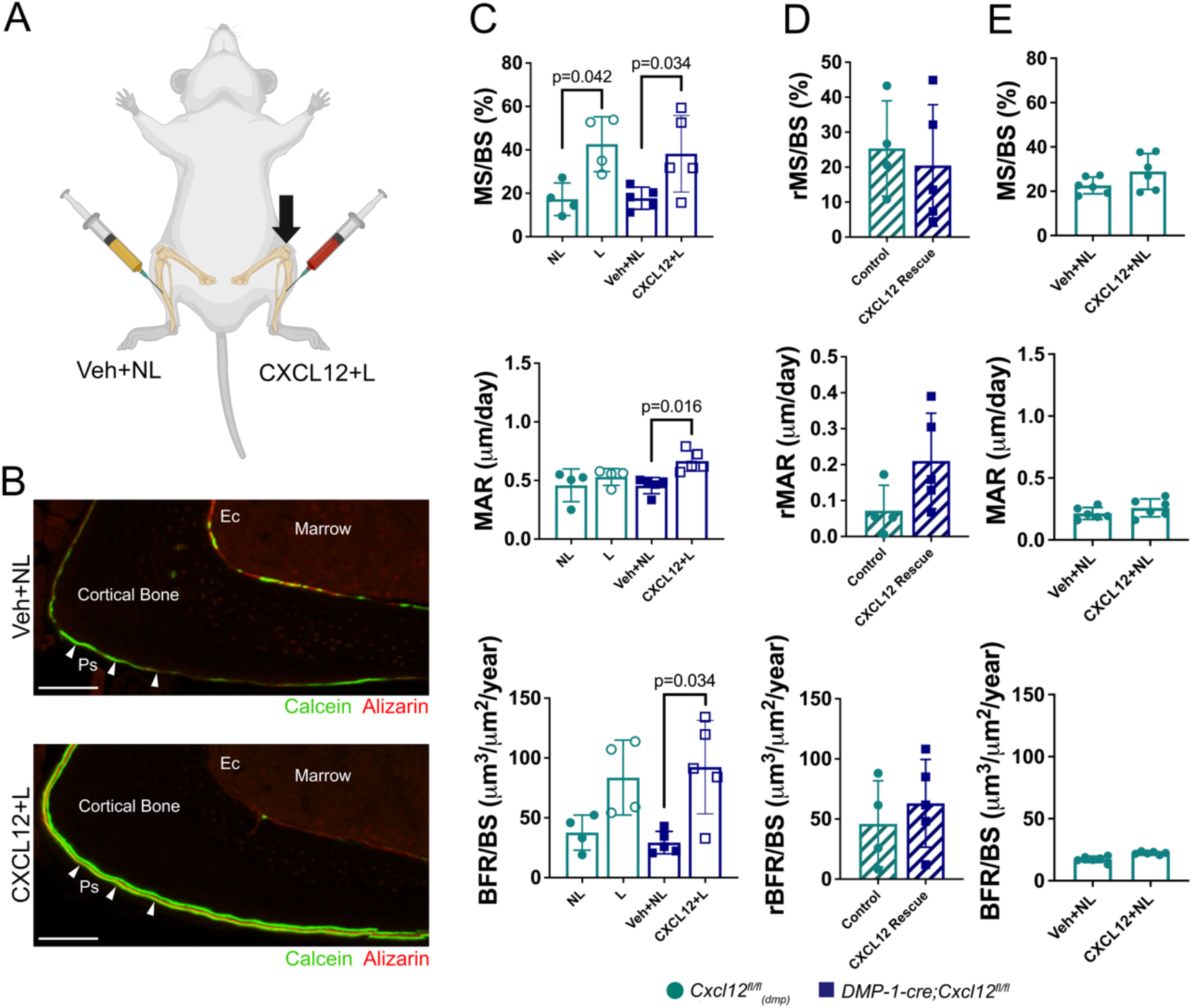
Exogenous CXCL12 treatment rescues the bone loading phenotype. (A) Experimental approach for rescuing load-induced bone formation using recombinant CXCL12 treatment. (B) Representative confocal images of a vehicle-treated and non-loaded (Vehicle+NL) tibia and a contralateral CXCL12-treated and loaded tibia from a *DMP-1-cre;Cxcl12^fl/fl^* mouse with calcein (green) and alizarin (red) fluorochrome bone labels on the periosteal (Ps, white arrows) and endocortical (Ec) surfaces. (C) Mineralizing surface (MS/BS), mineral apposition rate (MAR) and bone formation rate (BF/BS) in nontreated tibias from littermate control *Cxcl12^fl/fl^*_(*dmp*)_ mice (NL=non-loaded; L=loaded) and *DMP-1-cre;Cxcl12^fl/fl^* mice treated with vehicle (Veh+NL) or CXCL12 (CXCL12+L). (D) Relative bone formation parameters (loaded minus non-loaded) calculated from panel C. (E) Baseline bone formation parameters in non-loaded (NL) tibias from C57Bl/6 mice that were vehicle-treated (Veh+NL) or CXCL12-treated (CXCL12+NL). N ≥ 4. Data are presented as mean±SD.

### CXCL12/CXCR4 signaling regulates migration and osteogenic differentiation of bone marrow stromal cells

The finding that *Cxcl12* deletion in osteocytes and osteoblasts led to attenuated load-induced bone formation suggested that recruitment and/or differentiation of osteoprogenitors at the bone surface were dysregulated in the absence of CXCL12 signaling. Thus, we next investigated the role of the CXCL12/CXCR4 signaling axis in cell migration and osteogenic differentiation. Using a transwell migration assay, CXCL12 treatment at 10,000 ng/mL resulted in a significant increase in migration of primary bone marrow stromal cells (**Fig. 5A**). To determine whether CXCL12 signaling was an essential regulator of osteogenesis, primary bone marrow stromal cells from CXCL12 floxed mice were treated with Cre recombinase (CreR) to drive *cxcl12* gene deletion and cultured in osteogenic media. CreR treatment did not affect cell viability (**Fig. 5B**) as evaluated by an MTT assay. *Cxcl12* deletion led to a 37% decrease in CXCL12 expression and resulted in significantly lower expression of Runx2 (−67%) and Osx (−86%) (**Fig. 5C**) in BMSCs grown in osteogenic differentiation media. To demonstrate that CXCR4, the seven-pass G-protein-coupled receptor for CXCL12, was the target for CXCL12 in our system, we next showed that *cxcr4* gene deletion (−41%, **Fig. 5B**) resulted in significantly lower expression of Runx2 (−25%) and Osx (−43%) (**Fig. 5D**). Significantly lower expression of Osx was also observed following treatment with the CXCR4 antagonist AMD3100 compared to vehicle-treated controls (−57%) (**Fig. 5D**).

**Figure 5.**
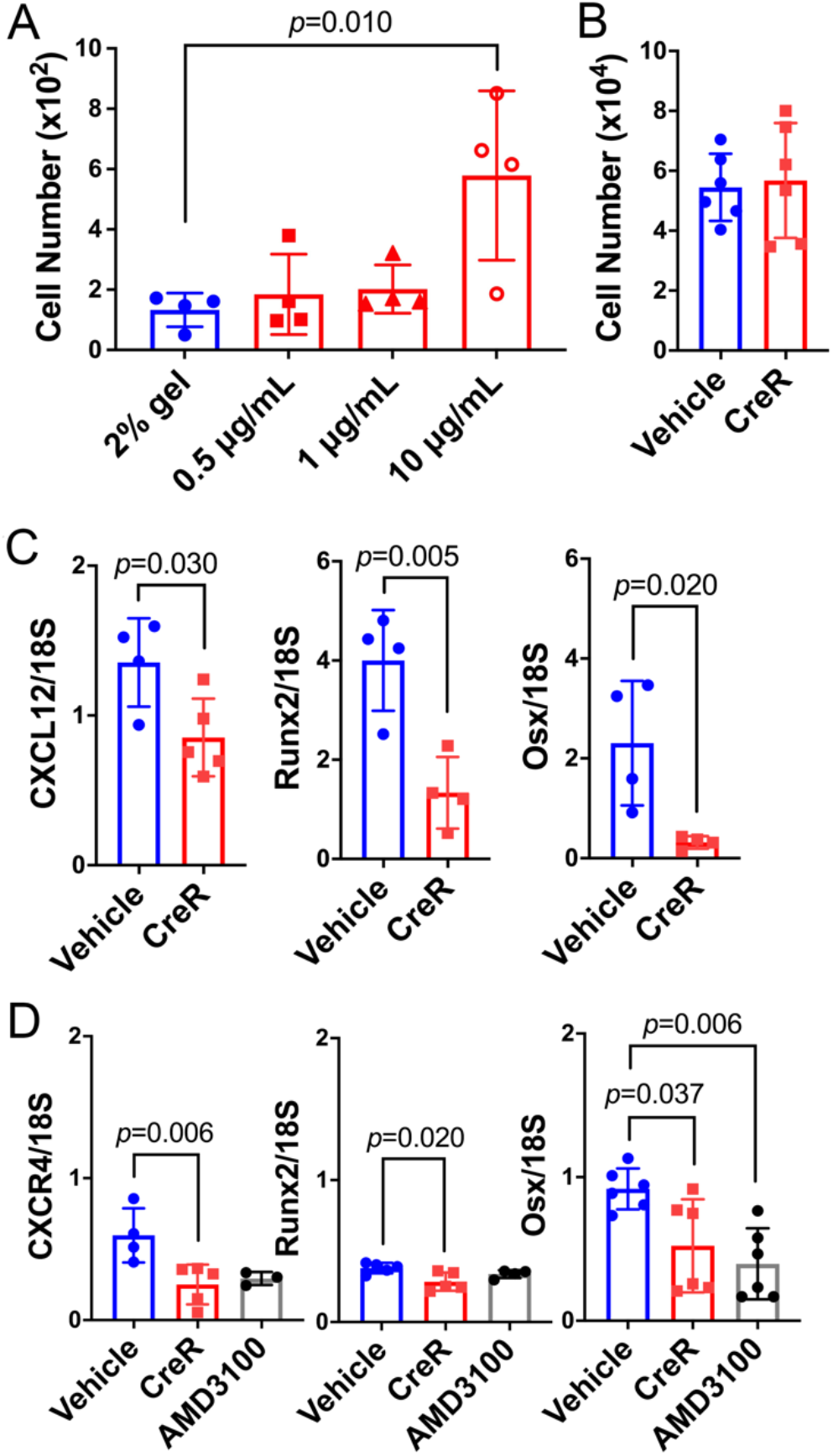
CXCL12/CXCR4 signaling enhances cellular migration and regulates osteogenic differentiation of mouse bone marrow stromal cells (mBMSCs). (A) Quantification of mBMSC migration after treatment with 0.5 μg/mL, 1 μg/mL and 10 μg/mL of recombinant CXCL12 protein. (B) Quantification of mBMSC number following treatment with Cre recombinase (CreR). (C) mRNA expression of CXCL12, Runx2 and Osx in mBMSCs isolated from *Cxcl12^fl/fl^* mice following CreR treatment. (D) mRNA expression of CXCR4, Runx2 and Osx in BMSCs isolated from *Cxcr4^fl/fl^* mice following treatment with CreR and AMD3100. N ≥ 4. Data are presented as mean±SD.

### CXCL12/CXCR4 signaling synergizes Wnt signaling pathway activation

The data described above together suggest that CXCL12, signaling via CXCR4, regulates osteogenic differentiation of bone marrow stromal cells. To identify a downstream signaling mechanism governing this response, we focused on the Wnt/β-catenin signaling pathway given its predominant role in osteogenesis (55–57). Wnt signaling results in cytoplasmic accumulation and nuclear translocation of β-catenin by inactivating, via phosphorylation, the enzyme glycogen synthase kinase-3β (GSK-3β), a member of the β-catenin destruction complex (58). We first showed that 1-hour treatment with CXCL12 alone and Wnt3a+CXCL12 resulted in a significant increase in GSK-3β phosphorylation compared to the vehicle control, an effect that was inhibited by cotreatment with the CXCR4 antagonist AMD3100 (**Fig. 6A,B**). We next showed an increase in β-catenin protein expression (**Fig. 6C,D**) and nuclear translocation (**Fig. 6E**) following a 4-hour co-treatment with Wnt3a+CXCL12. A separate experiment confirmed a significant increase in Axin2 gene expression, a Wnt/β-catenin target gene, following treatment with both Wnt3a and CXCL12 for 72 hours compared to each alone and to the vehicle control (**Fig. 6F**). Together, these data suggest that CXCL12/CXCR4 signaling can synergize Wnt signaling by phosphorylating the protein kinase GSK-3β leading to accumulation and nuclear translocation of β-catenin, and an upregulation Axin2, a Wnt target gene.

**Figure 6:**
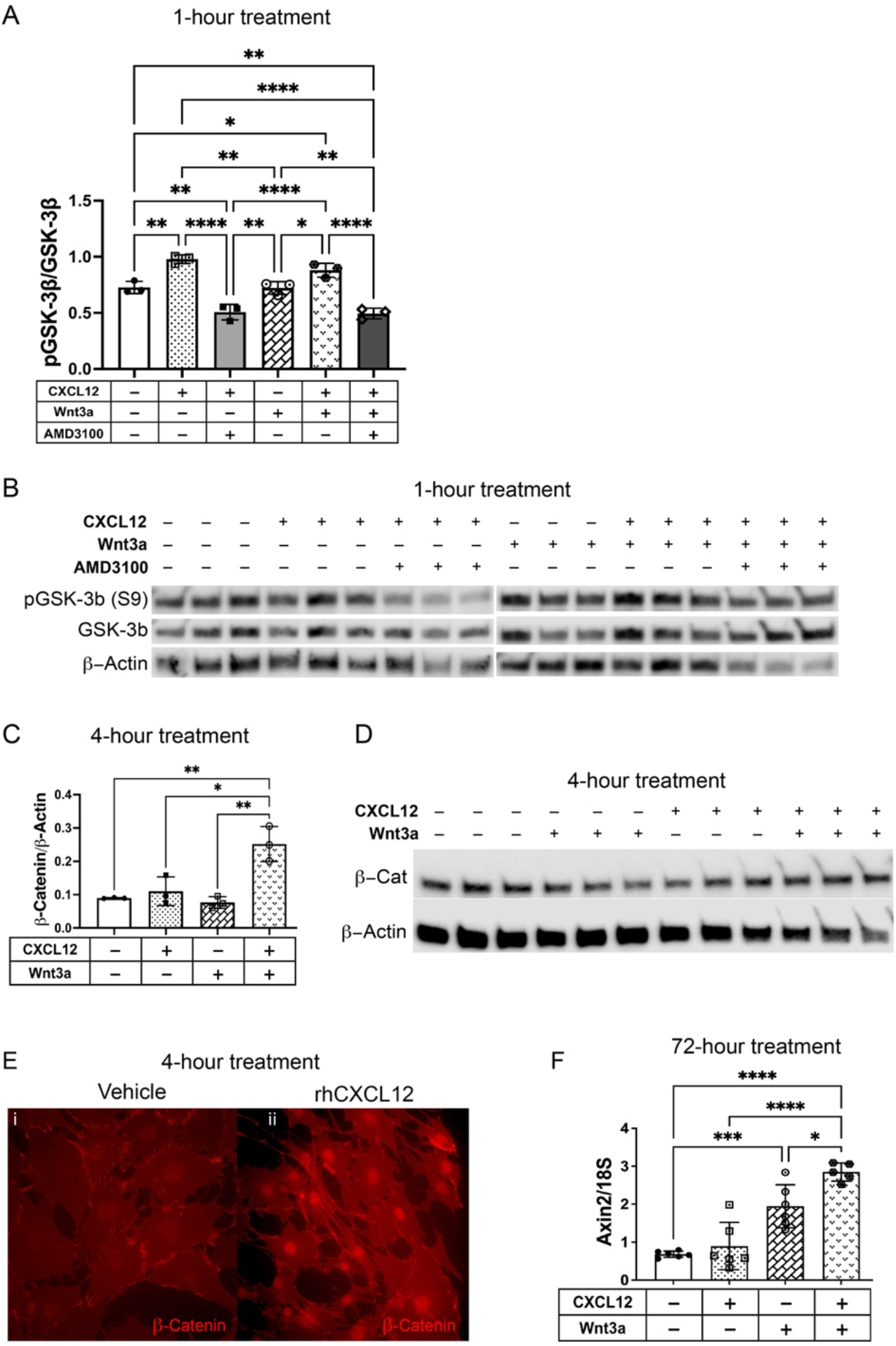
CXCL12 treatment synergizes with the Wnt signaling pathway. (A,B) Western blot of phosphorylated GSK-3β in BMSCs isolated from *Cxcl12^fl/fl^* mice following a 1-hour incubation in serum-free media containing rhCXCL12, rhWnt3a, and/or AMD3100. (C,D) Western blot of β-catenin in BMSCs isolated from *Cxcl12^fl/fl^* mice following a 4-hour incubation in serum-free media containing rhCXCL12 and/or rhWnt3a. (E) Fluorescent microscopy image showing β-catenin staining in MC3T3-E1 subclone 4 preosteoblasts following a 4-hour incubation with (i) vehicle and (ii) rhCXCL12. (F) qRT-PCR analysis of Axin2 at the mRNA level 72 hr after treatment with rhCXCL12 and rhWnt3a. N ≥ 3. *p-values:* * < 0.02, ** < 0.002, *** < 0.001, **** < 0.0001.

## DISCUSSION

The cellular and molecular events through which osteoanabolic pathways are triggered in response to mechanical loading in bone are incompletely understood. The current studies provide evidence that the CXCL12/CXCR4 signaling axis is essential for load-induced bone formation and synergizes the Wnt-β-catenin pathway to promote osteogenesis. C-X-C Motif Chemokine Ligand 12 (CXCL12), a chemokine involved in cell migration (44, 59), proliferation (36), survival and apoptosis (60), is highly expressed by skeletal stem and progenitor cells (44, 48), with decreasing expression in bone lineage cells as osteogenic differentiation progresses (36, 61). We previously demonstrated that *in vivo* bone anabolic mechanical loading of the mouse ulna results in increased expression of CXCL12 in osteocytes and bone lining cells in mouse cortical bone and that osteoblast-like cells grown in culture release CXCL12 in response to increased fluid flow shear stress (45). In the same study, we showed that systemic antagonism of CXCR4, the G protein-coupled receptor for CXCL12, attenuates load-induced bone formation. One limitation of that work was the inability to distinguish the cell-specific source of CXCL12 involved in regulating osteogenesis following loading, knowledge important for identifying treatments targeting load-induced bone formation.

In the present study, we report the effects of CXCL12+ cell ablation and *cxcl12* gene knockout on adult bone in homeostasis and following mechanical loading. We demonstrate that mice subjected to targeted ablation of CXCL12+ cells, and mice harboring osteocyte- and osteoblast-specific *cxcl12* gene knockdown, separately, fail to fully adapt to an applied whole bone mechanical stimulus. Given that collagen 1α1+ osteoblasts and DMP-1+ osteocytes appear during late-stage osteogenic lineage commitment (62, 63), our findings suggest a model in which mechanical loading leads to up-regulation and release of CXCL12 by osteoblasts and osteocytes (45), which then binds to its membrane-bound receptor CXCR4 expressed on SSPCs residing on bone surfaces, regulating their behavior and function. This paracrine signaling model is supported by our *in vitro* data showing that CXCL12 enhances migration of primary bone marrow stromal cells (BMSCs) and by previously published data showing that CXCL12 regulates BMP-2-induced osteogenic differentiation of BMSCs (48). Autocrine regulation via CXCL12 signaling likely also contributes to load-induced bone formation given that SSPCs express both CXCL12 (36) and its receptor CXCR4 (53). Indeed, our data show that *in vitro* deletion of *cxcl12* and *cxcr4*, separately, in primary BMSCs significantly attenuates their osteogenic differentiation. These results are supported by previous work showing that conditional *in vivo* deletion of *cxcl12* in Prrx-1+ and Osx+ cells, separately, leads to a significant decrease in trabecular bone volume and reduced expression of osteogenic markers in whole femur extracts (41).

The molecular mechanism by which CXCL12/CXCR4 signaling regulates load-induced bone formation is not well understood. Mechanical loading upregulates expression of CXCL12 (45) and Wnt (30, 64), suggesting these signals may cooperatively control osteogenesis. Our *in vitro* data demonstrate that CXCL12 treatment mediates the canonical Wnt pathway by increasing phosphorylation of GSK-3β, an important component of the β-catenin destruction complex and whose phosphorylation leads to its inactivation (65). This effect that was blocked by treatment with a CXCR4 antagonist. Co-regulation of the CXCL12/CXCR4 and Wnt signaling pathways has been demonstrated in other systems. CXCL12 signaling induces phosphorylation of GSK-3β in human dental pulp stem cells (66) and human THP-1-derived macrophages (67), which supports our findings. Conversely, Wnt can regulate CXCL12 expression in mammary gland cancer cells owing to three consecutive TCF/LEF consensus sequences in the *cxcl12* gene (68). Inhibition of the Wnt/β-catenin pathway in fibroblasts and HSCs leads to a complete inhibition of TGFβ-induced CXCL12 expression (69). These data together suggest a model by which loading upregulates CXCL12 expression in osteoblasts and osteocytes, which, in turn, synergizes the Wnt signaling pathway, by phosphorylating GSK-3β, to enhance osteogenic differentiation. Of note, CXCL12 has also been shown to regulate osteogenic differentiation by modulating BMP-2-induced Smads and ERK activation (48), which can, in turn, modulate CXCL12 expression (70–73); thus, CXCL12 appears to work in concert with other known osteogenic signaling pathways to regulate bone formation. Whether CXCL12 and BMP signaling are together regulating load-induced bone formation requires further study.

To further support a paracrine signaling model, exogenous treatment of CXCL12 in mechanically loaded *Col1α1-cre;Cxcl12^fl/fl^* animals rescued the diminished bone formation rate – a product of mineralizing surface and mineral apposition rate – to levels at or above their respective phenotype controls. This effect was due to an increase in both mineralizing surface and mineral apposition rate in treated tibiae, suggesting that both the number and bone-forming activity of individual osteoblasts were enhanced with CXCL12 treatment. Of note, CXCL12 treatment did not lead to an increase in basal bone formation rates in normally loaded tibiae, implying that increased mechanical loading – or another bone anabolic stimulus – is required to fully exploit the osteogenic nature of CXCL12. This fits with what we know about crosstalk with other osteogenic signaling pathways (Wnts, BMPs) and by data showing that fluid flow shear stress leads to an increase in the CXCR4 expression in bone marrow stromal cells (45), which would in essence prime SSPCs to respond to CXCL12. Together, these data suggest that treatment with CXCL12 would be most effective at inducing new bone formation when combined with exercise or administered during fracture repair.

One important question to address when considering development of CXCL12 as a bone anabolic agent is which SSPC populations differentiate into bone-forming osteoblasts as a result of increased mechanical loading. Previous work shows that Prrx1+ (74, 75) and Osx+ (76, 77) osteoprogenitors residing in the periosteum exhibit increased proliferation with mechanical loading, while ablation of Prrx1+ cells leads to significantly attenuated load-induced bone formation (75), the later suggesting that a subset of bone-forming osteoblasts originate from the Prrx1+ cell population following mechanical loading. In fact, *cxcr4* gene deletion in the more differentiated Osx+ cell population results in significant decreases in bone mass and bone formation rates (53), suggesting that Osx+ cells are an important target of CXCL12 in the context of osteogenesis. Additional studies aimed at identifying and characterizing the different SSPC populations residing in the periosteum and their response to CXCL12/CXCR4 signaling in the context of mechanical loading are underway.

The fact that *cxcl12* deletion results in significantly lower trabecular, but not cortical, bone volume in female *Col1α1-cre;Cxcl12^fl/fl^* and *DMP-1-cre;Cxcl12^fl/fl^* mice is not surprising given the high turnover rate of trabecular bone and its dependence on the generation of new osteoblasts. Of note, our histological data did not show a significant decrease in osteoblast perimeter in 16-week-old mice, implying effects of *cxcl12* knockout on osteoblastogenesis may have appeared at an earlier age and persisted into adulthood. CXCL12 also regulates the maintenance of hematopoietic cells (46), from which osteoclasts are derived (78), including the release of hematopoietic cells from their niche (38, 79), as well as promoting the recruitment (80), development and survival of osteoclasts (60). Thus, in *Col1α1-cre;Cxcl12^fl/fl^* and *DMP-1-cre;Cxcl12^fl/fl^* mice, one might expect a decrease in osteoclast number, which we did not observe, lending credence to that idea that effects on bone volume developed during growth in these non-inducible mouse lines. Finally, it is unclear why lower trabecular bone volume was not observed in male mice, though there appears to be link between estrogen and CXCL12 (81, 82), a connection being clarified in separate studies.

Cre-lox recombination technology was used to generate two mouse lines in which CXCL12 was conditionally deleted in osteoblasts – and consequently in the downstream osteocyte phenotype – using the 2.3 kb Col1α1 promoter (83), and in primarily osteocytes, using the 9.6 kb DMP-1 promoter (63). The recombination efficiency assaysin the current study showed a 75% and 60% reduction in CXCL12 transcripts using the Col1α1 and DMP-1 promoters, respectively, similar to other gene ablation studies using Cre-lox recombination technology (63, 84). We specifically targeted osteocytes given their presumed role as primary mechanosensors and regulators of mechanically-induced bone formation (8, 14); that is, we hypothesized that release of CXCL12 from osteocytes in mechanically loaded bone is an important signaling event in the anabolic response. We also targeted mature osteoblasts for gene deletion to examine the relative effect of osteoblast-specific CXCL12 expression on skeletal homeostasis and mechanically-induced bone formation, recognizing that its expression may influence osteoblast function through autocrine signaling and/or influence the function of nearby cells, including osteoprogenitors, through paracrine signaling. This approach was somewhat limited given that DMP-1 is also expressed in some late-stage osteoblasts (63, 84, 85), bone lining cells (85), and intracortical perivascular cells (86). Indeed, using the Ai14 reporter mouse, we showed similar expression patterns of Col1α1 and DMP-1 in cortical bone and in the periosteum demonstrating some overlap of these two markers, which limited our ability to decisively show an osteocyte-versus osteoblast-driven effect.

The finding that osteoblast- and osteocyte-expressed CXCL12 regulates the anabolic response of bone to mechanical loading has important implications for the prevention and treatment of disuse and age-related (64, 74, 87–91) bone loss in humans. First, it suggests that the CXCL12/CXCR4 signaling axis may be a feasible target for promoting osteogenesis and increasing bone mass. Second, it indicates that bone anabolic exercise or co-treatment with Wnts or BMPs may be necessary to benefit from a CXCL12-based treatment. Whether CXCL12 treatment can enhance delayed fracture repair due to aging or traumatic volumetric bone loss is currently under investigation.

## MATERIALS AND METHODS

### Animals

All procedures were approved by the NYU Institutional Animal Care and Use Committee. *Cxcl12^fl/fl^* mice, which are commercially available (Jackson Laboratory, stock no. 022457), were crossed with 2.3kb Col1α1-Cre mice, which express Cre recombinase driven the 2.3kb murine *Col1a1* promoter (83), to generate CXCL12^flox/flox^;2.3kb Col1α1-Cre (*Col1α1-cre;Cxcl12^fl/fl^*) mice. *Cxcl12^fl/fl^* were also crossed with 10kb DMP-1-Cre mice, which express Cre recombinase driven by the 9.6-kb murine *Dmp1* promoter (63) to generate CXCL12^flox/flox^; 10kb DMP-1-Cre (*DMP-1-cre;Cxcl12^fl/fl^*) mice. Age-matched and sex-matched littermates were used as controls. *Cxcl12* deletion was confirmed by genomic DNA analysis (**Fig. S6A**), RT-PCR (**Fig. S6B**), protein blot (**Fig. S6C**) and immunohistochemistry (**Fig. S6D,E**). Genotype was determined using agarose gel electrophoresis of control and floxed *cxcl12*, Cre and 18s PCR products from genomic DNA. RNA was isolated from kidney, marrow and cortical bone, and expression of CXCL12 and 18S mRNA assessed using Taqman assays. CXCL12 expression in cortical bone, marrow and kidney was assessed by protein blot analyses (n=3/genotype) using recombinant human CXCL12 as a control (R&D, #350-NS).

Protein bands of interest were visualized using an ECL substrate kit (Bio-Rad Laboratories, #1705060S) and captured using an ImageQuant LAS 4000 imager. Protein expression was normalized to GAPDH (1:2500, Abcam, #ab9485). Each mouse line was crossed with the Ai14 reporter mouse (Jackson Laboratory, stock no. 007908), which harbors a *loxP*-flanked STOP cassette upstream of a CAG promoter-driven red fluorescent protein variant (tdTomato), to determine location of Col1α1 and DMP-1 expressing cells in mouse tibiae and to confirm osteoblasts and osteocytes as targets for gene deletion. Bones (n=3/genotype) were harvested, cryo-processed, mounted and imaged for DAPI and tdTomato using a Zeiss LSM710 confocal microscope with a 20× water immersion objective W Plan-Apochromat 20x/1.0 DIC M27 75mm (N.A. 1.0). See **Table S3** for primer sequences, Taqman assays and antibodies used for protein blots and immunohistochemistry. Body weight was assessed every two weeks from 4 to 16 weeks of age and no differences were observed (**Fig. S7**). Tibiae from 16- to 19-week-old female and male mice were used for experimentation with a sample size of *n*=6-15 depending on the outcome measure. *Cxcl12^tm2^.^1Sjm^/J* mice were obtained from Jackson Labs (stock #022458) (38) and express DsRedE2 from the endogenous *cxcl12* (chemokine (C-X-C motif) ligand 12) mouse promoter.

### Micro-computed x-ray tomography (μCT)

Tibiae (n=12-15/sex/genotype) were harvested, excised of soft tissue and stored in 70% ethanol for at least 72 hours prior to scanning. μCT was used to determine trabecular and cortical bone microarchitecture using a Bruker Skyscan 1172 with a 0.5 mm aluminum filter, at 56 kV, 179 μA, and exposure of 1850 ms and 6 μm^3^ voxel size. CT Analyzer (v1.16.4.1) was used to assess 3D microarchitecture in trabecular bone and 2D geometry in cortical bone. For trabecular bone, a volume of interest situated 250 μm distal from the end of the tibial growth plate and extending 1.5 mm towards midshaft was analyzed for trabecular bone volume fraction (BV/TV, %), thickness (Tb.Th, μm), number (Tb.N, 1/mm) and separation (Tb.Sp, μm). For cortical bone, a series of transverse slices spanning a 240 μm segment centered at the tibial mid-length was analyzed for total area (Tt.Ar, mm^2^), cortical area (Ct.Ar, mm^2^), thickness (Ct.Th, μm) and moments of inertia (Imax, Imin, mm^4^). Standard nomenclature was used for all measurements (54, 92). (n=10-14)

### Histology

Tibiae (n=4-5/sex/genotype) were harvested, fixed in 4% PFA at 4°C overnight and paraffin embedded. Longitudinal 5-μm thick sections were stained for tartrate-resistant acid phosphatase (TRAP) (93) and alkaline phosphatase (ALP) (94), dehydrated, mounted and imaged with an epifluorescent microscope. For each bone, a region of interest (ROI) including trabecular bone and marrow, which corresponded with the volume examined by μCT, was analyzed. The ROI was identified by tracing a line just below the growth plate, at the endocortical surfaces and extending down 1.75 mm away from the growth plate. For osteoblast activity, ALP-stained sections were analyzed for positively stained bone perimeter (Ob.Pm) normalized to total bone (B.Pm). For osteoclast number, TRAP-stained sections were analyzed for the total number of osteoclasts (N.Oc), the number of osteoclasts per bone surface (N.Oc/B.Pm), and the number of osteoclasts located in the marrow (N.Oc) – not associated with a bone surface.

### Biomechanical testing

Biomechanical properties of the femur were assessed using three-point bending (54). Femurs from female and male mice (n=6/sex/genotype) were harvested, wrapped in saline-soaked gauze and stored at −20°C prior to mechanical testing for at least 72 hours. A Bose Enduratec 3200 MTS vertical mechanical testing system with a 50 N load cell and a 7 mm span of the femur was used to determine whole bone stiffness, peak force, yield force and post yield displacement using a custom Matlab routine. Microarchitecture of the mid-femoral region was used to estimate material level properties including elastic modulus, yield stress and ultimate stress.

### Dynamic histomorphometry

Baseline bone formation rates were assessed in *Col1α1-cre;Cxcl12^fl/fl^, DMP-1-cre;Cxcl12^fl/fl^* mice and their WT littermates to test the effect of CXCL12 deletion in the developing skeleton. Beginning at 4 weeks of age, mice (n=12-15/sex/genotype) were injected with alternating fluorochrome bone labels (alizarin red, calcein and tetracycline) every 2 weeks. Tibiae were harvested two days after the last injection at 16 weeks of age, embedded in MMA, and sectioned with a diamond saw prior to imaging with the Zeiss LSM710 confocal microscope described above. Standard static and dynamic bone formation parameters 8/25/22 10:43:00 AM were determined at mid-length of each tibia using ImageJ during two periods spanning 4-8 and 10-14 weeks of age, separately. Static morphometric measurements included bone surface perimeter (B.Pm, μm), single and double labeled surface perimeter (sL.Pm, dL.Pm, μm, respectively), and interlabel area (Ir.L.Ar, μm). Calculated dynamic bone formation parameters included mineralizing surface (MS/BS = 100*[0.5*sL.Pm+dL.Pm] /B.Pm, %), mineral apposition rate (MAR = Ir.L.Ar /dL.Pm/days between labels, μm/day) and bone formation rate (BFR/BS =MAR*[MS/BS]*3.65, μm^3^/μm^2^/year).

### *In vivo* mechanical loading

The anabolic response to exogenous mechanical loading was assessed using the tibial axial compression model following a load-strain calibration procedure described previously (95). Sixteen-week-old *Cxcl12-creER;R26R^tdTomato^* and *Cxcl12-creER;Rosa26^lsl-tdTomato/iDTA^* mice were injected with tamoxifen (50 mg/kg in corn oil) 7 days prior to the first bout of loading to ablate CXCL12+ cells. Right tibiae from *Cxcl12-creER;R26R^tdTomato^, Cxcl12-creER;Rosa26^lsl-tdTomato/iDTA^, Col1α1-cre;Cxcl12^fl/fl^, DMP-1-cre;Cxcl12^fl/fl^*, and their WT littermates (*n*=5-10/sex/genotype) were loaded, with the left limb serving as the non-loaded control. The loading protocol consisted of a cyclic haversine axial compressive force (6 N peak load, 2 Hz, 120 cycles, 3X per week for 2 weeks) which generated 1100 με peak-to-peak strains at the periosteal surface. Animals were injected with alternating calcein and alizarin red fluorochrome bone labels on days 5, 12 and 19, respectively, and euthanized on day 21. Tibiae were harvested, embedded in MMA and sectioned for analysis of bone formation rates, as described above, between days 5-12, and days 12-19, after the first day of loading. In addition, dsRed-CXCL12 mice (n=2) were used to localize CXCL12 expression following mechanical loading, where right tibiae were loaded daily for four days, and bones harvested and subsequently processed for thick cryosection tissue imaging described above.

### *In vivo* administration of CXCL12

To determine whether CXCL12 treatment could rescue the attenuated load-induced osteogenesis phenotype in *DMP-1-cre;Cxcl12^fl/fl^* mice, they and their controls were treated with recombinant human CXCL12 (R&D Systems, #350-NS) in a biopolymer delivery vehicle (CellMate3D, BRTI Life Sciences, #CM-1002) immediately following mechanical loading. Recombinant human CXCL12 was reconstituted to a concentration of 100 μg/mL with sterile PBS per the manufacturer’s recommendation, followed by reconstitution of CellMate3D with the CXCL12 or PBS control solution. A final volume of 30 μL containing 1 μg CXCL12 was injected under the dermis of the anteromedial aspect of the loaded right tibia at mid-length using a 21-gauge needle on days 1 and 8, while the left non-loaded controls were injected with vehicle only. A separate group of WT littermates were used as non-loaded CellMate3D controls (n=4), whereby left tibiae were injected with CellMate3D vehicle only, and the right tibiae were treated with vehicle and CXCL12.

### Migration assay

MC3T3 cells were cultured in growth media and treated with increasing concentrations of recombinant human CXCL12/SDF-1α (R&D Systems, #350-NS) reconstituted in PBS and 0.1% BSA in a 24-well trans-well plate and allowed to migrate for 3 hours prior to cell quantification.

### *In vitro* osteogenic differentiation

To elucidate the mechanism underlying the attenuated *in vivo* bone loading response in *Col1α1-cre;Cxcl12^fl/fl^* and *DMP-1-cre;Cxcl12^fl/fl^* mice, the role of CXCL12 signaling in *in vitro* osteogenic differentiation was evaluated. Mouse bone marrow stromal cells (mBMSCs) were isolated as described previously (96, 97). Briefly, after euthanasia, tibial and femoral bone marrow from *Cxcl12^fl/fl^* and *Cxcr4^fl/fl^* mice, separately (n=3-4 per genotype), were flushed out with growth media containing DMEM (ATCC, #30-2002), 10% FBS (ATCC, #30-2020) and 1% Streptomycin/Penicillin (Fisher Scientific, #15140163) using a 3 ml syringe and 21G needle, into a 50 ml falcon tube under sterile conditions. Cells were plated in a 75 ml flask and incubated in a humidified chamber at 37°C with 5% CO_2_ for 5 days without feeding. Cells were washed with DPBS (PromoCell, #C40232), split into two 75 ml flasks and fed with growth media until 80-85% confluent. One flask of cells was treated with 1-2 μM (98, 99) TAT-Cre Recombinase (EMD Millipore, #SCR508) while the other was treated with DPBS alone. Cell viability following Cre-recombinase treatment was assessed using an MTT proliferation assay (ATCC, #30-1010K), separately. After 24 hours, flasks were washed with DPBS twice and cells fed with growth media until flasks were 95% confluent. Cells from each flask were then trypsinized and seeded into 6-well dishes at 5 × 10^4^ cells/well for a total of 8 days. Cells were fed with either growth media or StemXVivo osteogenic media (R&D Systems, #CCM007 and CCM009). In addition, to determine whether signaling via CXCR4 is important for osteogenic differentiation, *Cxcr4^fl/fl^* BMSCs were also treated with AMD3100 (200 ng/ml, Sigma-Aldrich, #A5602), a CXCR4 antagonist. Treated cells were rinsed with PBS and lysed using 600 μl of lysis RLT lysis buffer per manufacturer protocol (Qiagen, #74134). Total mRNA was reverse-transcribed to cDNA using the Omniscript Reverse Transcriptase kit (Qiagen, #205111). TaqMan gene expression master mix (Fisher Scientific, #43-690-16) and assays (Table S3) were used for real-time qPCR (Thermo Fisher, Applied Biosystems QuantStudio 3 and BioRad CFX 384) to determine CXCL12, CXCR4, Runx2 and Osx expression. 18S was used as the housekeeping gene.

### CXCL12 and Wnt signaling crosstalk

We next tested our hypothesis that CXCL12 is acting via the Wnt signaling pathway to regulate osteogenesis. First, primary BMSCs isolated from C57BL/6 mice were plated in 6-well dishes until 80% confluent and serum starved (0.5% FBS) for 48 hours prior to treatment with one or a combination of CXCL12 (200 ng/ml, R&D Systems, 460-SD), Wnt3a (100 ng/ml, R&D Systems, 1324-WN-002/CF) and AMD3100 (200 ng/ml, Sigma-Aldrich, 239820). Treated cells were incubated for 1 and 4 hours and total protein was isolated using RIPA lysis and extraction buffer (Thermo Scientific, 89900) and run using SDS-PAGE in 10% precast polyacrylamide gels (BioRad, 1620145) and transferred to nitrocellulose membranes (BioRad, 1620145). After blocking in 5% non-fat dry milk in TBST, GSK3b, pGSK3b and β-catenin were detected using primary antibodies and an HRP-linked secondary antibody, followed by ECL visualization. β-Actin was used as the housekeeping protein. Next, MC3T3-E1 Subclone 4 preosteoblasts (ATCC CRL-2593) grown in monolayer were treated with vehicle or CXCL12 (20 μg/ml, R&D Systems, 460-SD) for 4 hours. Cells were then fixed in 4% paraformaldehyde and permeabilized with 0.1% Triton-X in PBS. Cells were then incubated with 1:200 monoclonal mouse anti-β-catenin antibody and 1:100 donkey IgG in primary blocking solution (PBS, 1% BSA, 0.1% NP-40) overnight. Cells were washed 3 x 10 minutes in PBS and incubated with goat anti-mouse Alexa Fluor 594 for 4 hours. Cells were imaged with the EVOS FL Auto light microscope (Thermo Fisher, 20X Obj, AMEP4653 470/22 excitation and 510/42 emission filter) and assessed for β-catenin translocation to the nucleus. In an experiment to determine activation of the Wnt signaling pathway in response to CXCL12 signaling, primary BMSCs were plated in 6-well dishes until 80% confluent and serum starved (0.5% FBS) for 72 hours prior to treatment with one or a combination of CXCL12 (200 ng/ml, R&D Systems, 460-SD) and Wnt3a (100 ng/ml, R&D Systems, 1324-WN-002/CF). Treated cells were incubated for 48 hours and Axin2 gene expression was evaluated using the protocol described above. All primers, probes and antibodies are listed in Table S3.

### Flow Cytometry

After removing distal epiphyseal growth plates and cutting off proximal ends, femurs were cut roughly and incubated with 2 Wunsch units of LiberaseTM (Sigma/Roche 5401127001) and 1 mg of Pronase (Sigma/Roche 10165921001) in 2 ml Ca2+, Mg2+-free Hank’s Balanced Salt Solution (HBSS, Sigma H6648) at 37 °C for 60 min. on a shaking incubator (ThermomixerR, Eppendorf). After cell dissociation, cells were mechanically triturated using an 18-gauge needle with a 1 ml Luer-Lok syringe (BD) and a pestle with a mortar (Thermo Fischer), and subsequently filtered through a 70 μm cell strainer (Falcon) into a 50 ml tube on ice to prepare single cell suspension. These steps were repeated three times. Dissociated cells were stained and sorted to enrich for the stromal cell populations for 30 minutes on ice. Flow cytometry analysis was performed using a ZE5 Yeti Analyzer. Acquired raw data were further analyzed on FlowJo software (TreeStar).

### Data analyses

Data are reported as mean ± SD. Prism 6 Statistical Software (GraphPad Software, Inc.) was used for all analyses. Differences in the slopes and intercepts of the load-strain calibration curves were tested using linear regression analysis. Differences in trabecular bone microarchitecture parameters, cortical bone geometric properties, osteoclast and osteoblast numbers, and bone formation parameters were tested for significance using a repeated measures two-way ANOVA with posthoc Tukey’s correction for multiple comparisons with genotype and knockdown line, or loading and treatment as the main factors. In vitro cell migration and gene expression data were tested using an ordinary one-way ANOVA with Tukey’s multiple comparison. Significance for all tests was set at α=0.05 and β≤0.20.

## Acknowledgements

μCT scanning was performed in the NYU Tandon Maker Space. Microscopy was performed in the NYU Langone Microscopy Laboratory. This work was supported in part by a VA Career Development Award (ABC), a VA Merit Review Award 1I01RX001500 (ABC), and NIH Grant AR073864 (ABC) and an NYU Clinical and Translational Science Institute Postdoctoral Fellowship 5TL1TR001447-03 (PCZ). The content is solely the responsibility of the authors.

## Notes

### Competing Interest Statement

The authors have declared no competing interest.

