## Supplemental Information for "CXCL12 in late-stage osteoblasts and osteocytes is required for load-induced bone formation in mice"

**This PDF file includes:**

Supporting text
Figures S1 to S7
Tables S1 to S3
Legends for Movies S1 to S2

**Other supporting materials for this manuscript include the following:**

Movies S1 to S2

**Supporting Information**

**SI Figures**

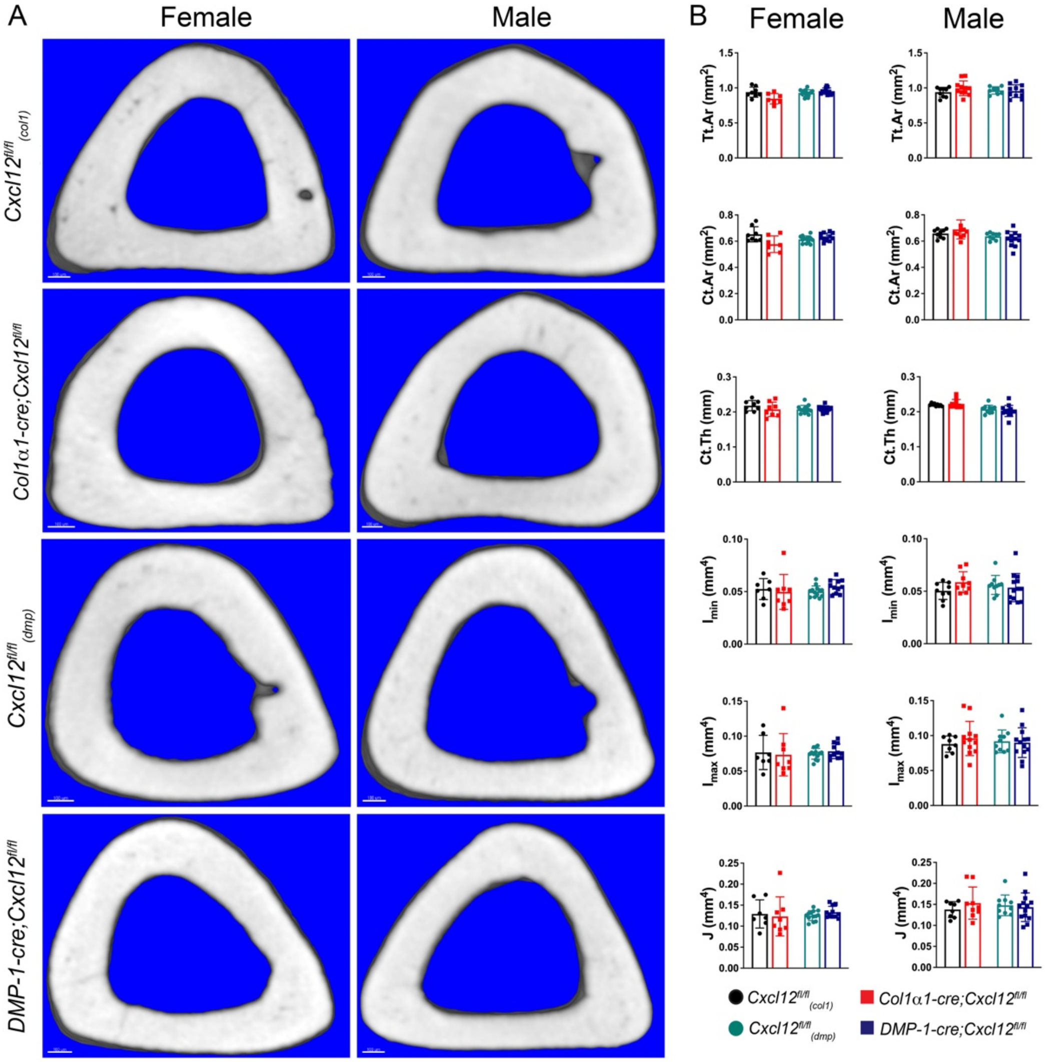

**Fig. S1.** Cortical bone structure *Col1α1-cre;Cxcl12$^{fl/fl}$* and *DMP-1-cre;Cxcl12$^{fl/fl}$* mice. (A) Representative μCT renderings of cortical bone at tibial midshaft in transverse view from 16-week-old female and male *Col1α1-cre;Cxcl12$^{fl/fl}$* and *DMP-1-cre;Cxcl12$^{fl/fl}$* mice and their respective controls. Scale bar, 100 μm, n ≥ 7. (B) Quantification of tissue area (Tt.Ar) (bone and marrow), cortical bone area (Ct.Ar), cortical bone thickness (Ct.Th), minimum second moment of area (Imin), maximum second moment of area (Imax) and polar moment (J). Data are presented as mean±SD.

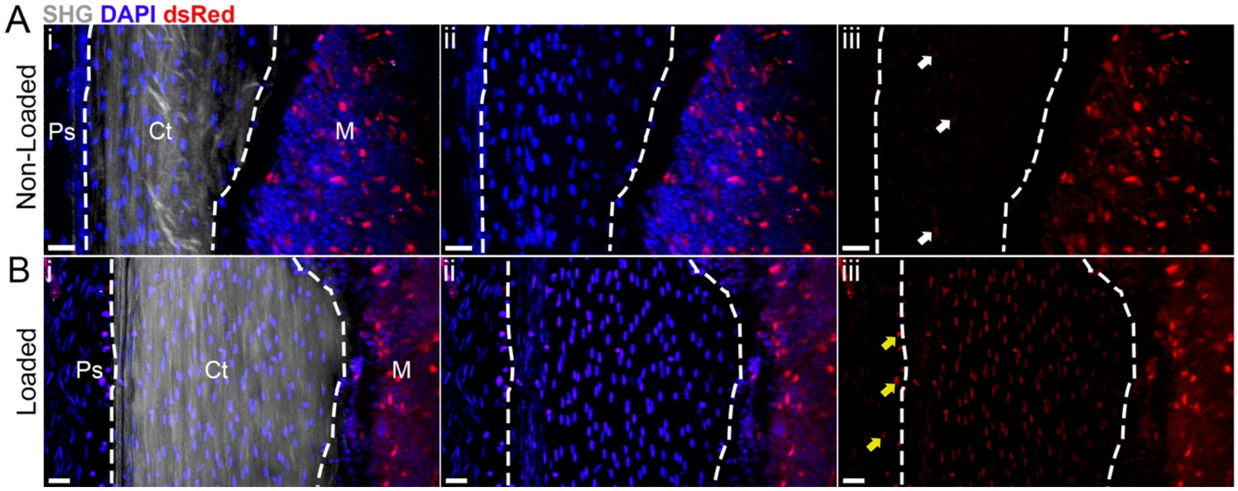

**Fig. S2.** CXCL12 expression in non-loaded and loaded mouse tibiae. Representative longitudinal confocal and 2-photon images of (A) non-loaded and (B) loaded tibiae at mid-shaft from CXCL12-dsRed mice showing periosteum (Ps), cortical bone (Ct), and marrow (M). (Ai, Bi) second harmonic generation (SHG, gray) signal indicates cortical bone (Cb) outlined in white dotted lines and adjacent to the periosteum (Ps) on the left and bone marrow (M) on the right. Overlay: (Ai, Bi) SHG (gray), DAPI (blue) and dsRed (red) expression; (Aii, Bii) DAPI and dsRed expression only, and (Aiii, Biii) dsRed expression only. (Ai,ii,iii) In non-loaded bone, dsRed expression is observed primarily in the marrow and in some cortical bone osteocytes (white arrows) (Aiii). (Bi,ii,iii) In loaded bone, dsRed expression is observed in the marrow, (Biii) cortical bone osteocytes and in cells in the periosteum (yellow arrows). Scale bar = 20 µm.

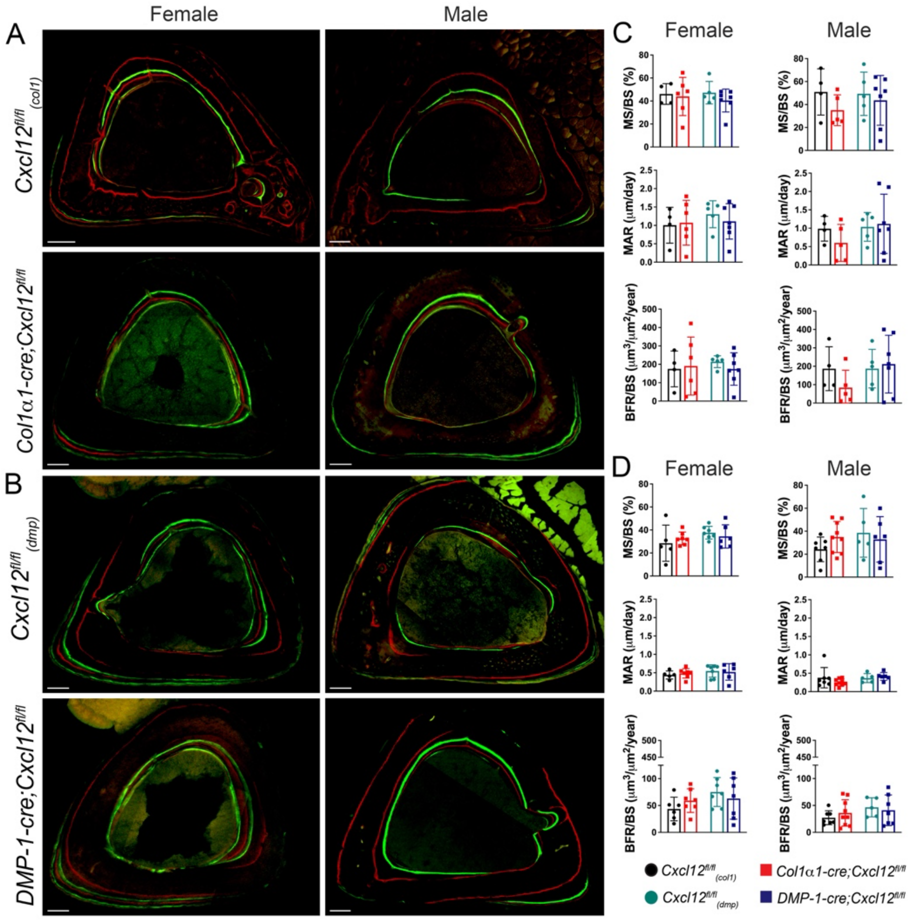

**Fig. S3.** Basal bone formation rates in the tibia from *Col1α1-cre;Cxcl12*$^{fl/fl}$ and *DMP-1-cre;Cxcl12*$^{fl/fl}$ mice. Representative images of the tibial midshaft in transverse section from female and male (A) *Col1α1-cre;Cxcl12*$^{fl/fl}$ and (B) *DMP-1-cre;Cxcl12*$^{fl/fl}$ mice and their respective controls, *Cxcl12*$^{fl/fl}_{(col1)}$ and *Cxcl12*$^{fl/fl}_{(dmp)}$. Mice received alternating alizarin red injections at 4, 10 and 16 weeks of age and calcein (green) injections at 8 and 14 weeks of age. Mineralizing surface (MS/BS), mineral apposition rate (MAR) and bone formation rates (BFR/BS) between (C) 4 and 8 weeks of age and (D) 10 and 14 weeks of age. N ≥ 5.

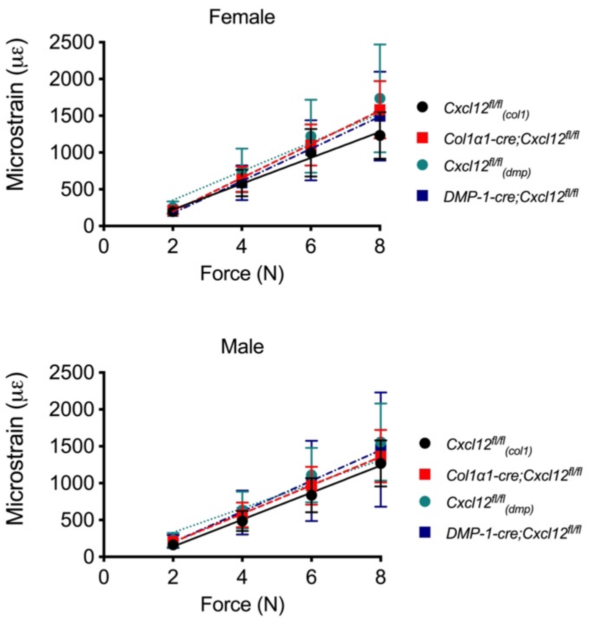

**Fig. S4.** Load-strain calibration curves for mouse tibiae. Load-strain curves for tibiae from female and male *Col1α1-cre;Cxcl12*$^{fl/fl}$ and *DMP-1-cre;Cxcl12*$^{fl/fl}$ mice and their respective controls subjected to cyclic axial compressive loading were generated by plotting sequential increasing peak loads and the resulting peak microstrain. N ≥ 6.

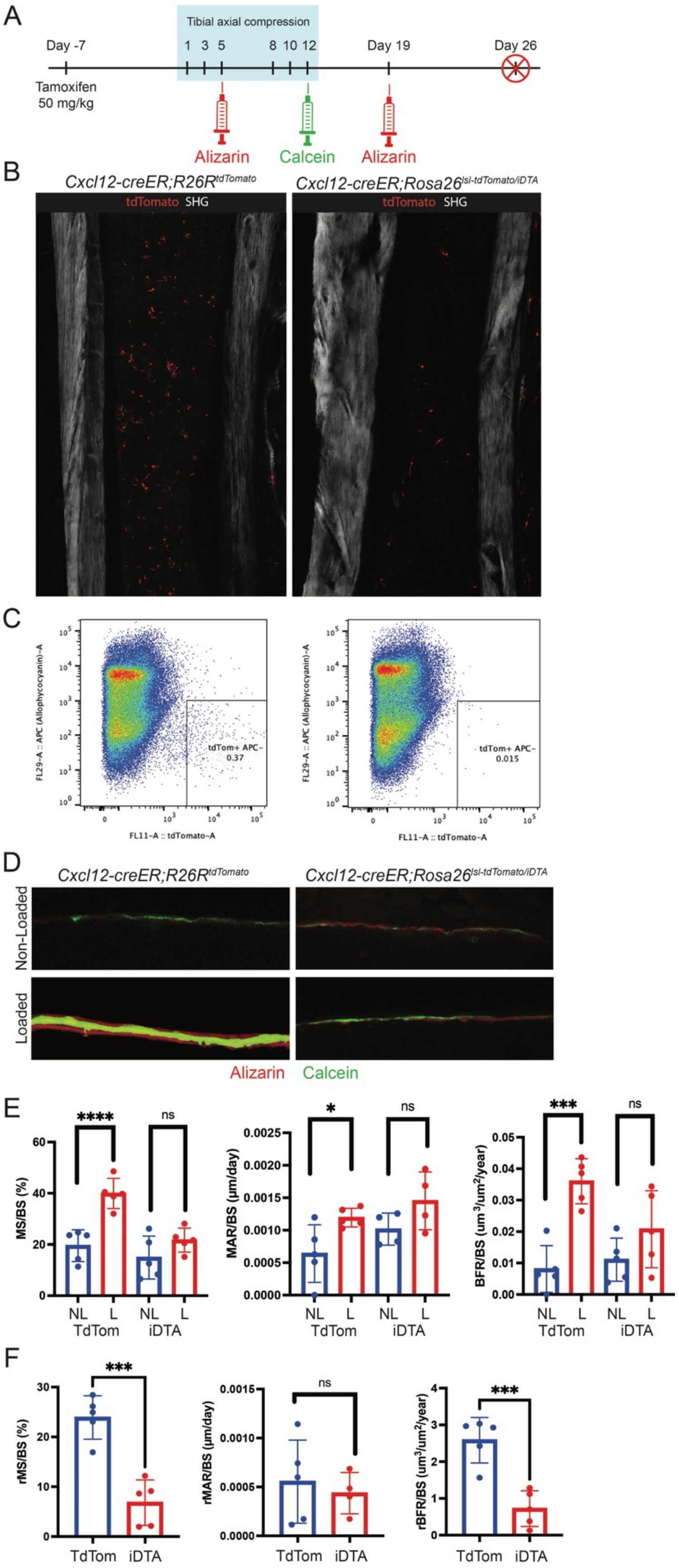
A
Tibial axial compression
Day -7
1 3 5 8 10 12
Day 19
Day 26
Tamoxifen
50 mg/kg
Alizarin
Calcein
Alizarin
B
Cxcl12-creER;R26R tdTomato
Cxcl12-creER;Rosa26 lsl-tdTomato/iDTA
tdTomato SHG
tdTomato SHG
C
FL29-A :: APC (Allophycocyanin)-A
FL11-A :: tdTomato-A
tdTom+ APC-
0.37
tdTom+ APC-
0.015
D
Cxcl12-creER;R26R tdTomato
Cxcl12-creER;Rosa26 lsl-tdTomato/iDTA
Non-Loaded
Loaded
Alizarin
Calcein
E
MS/BS (%)
MAR/BS (µm/day)
BFR/BS (um3/um2/year)
****
ns
*
ns
***
ns
NL L NL L
TdTom iDTA
F
rMS/BS (%)
rMAR/BS (µm/day)
rBFR/BS (um3/um2/year)
***
ns
***
TdTom iDTA

**Fig. S5.** CXCL12+ cells are essential for periosteal load-induced bone formation. (A) Schematic representation of the experimental timeline. Ablation was induced with a single tamoxifen dose (50 mg/kg) followed by a 7-day clearance period. Tibias were loaded on six occasions over a two-week period. Mineralizing surfaces were labeled with alizarin and calcein on days 5, 12 and 19. Mice were euthanized on day 26 for histological evaluation. (B) Tile-scan confocal images of 16-week-old *Cxcl12-creER;R26Rt*$^{dTomato}$ (control) and *Cxcl12-creER;Rosa26*$^{lsl\text{-}tdTomato/iDTA}$ (ablated) tibias in longitudinal section after a 7-day clearance period. CXCL12+ cells are shown in red. (C) Flow-cytometry analysis of FSC/SSC-gated cells isolated from whole bone. Left panel: scatter plot of cells from mice without tamoxifen recombination. Right panel: scatter plot of cells from mice with tamoxifen recombination after a 7-day clearance period. (D) Representative images of fluorochrome-labeled non-loaded (NL) and loaded (L) tibias illustrating periosteal bone formation in control and ablated mice. (E-F) Mineralizing surface (MS/BS), mineral apposition rate (MAR/BS) and bone formation rates (BFR/BS) in NL and L tibias from control (TdTom) and ablated (iDTA) mice. (F) Relative (r) (loaded minus non-loaded) mineralizing surface (rMS/BS), mineral apposition rate (rMAR/BS) and relative bone formation rates (rBFR/MS) in control (TdTom) and ablated (iDTA) mice. Data shown as mean ± SD. *$p < 0.05$ by a Student's t-test.

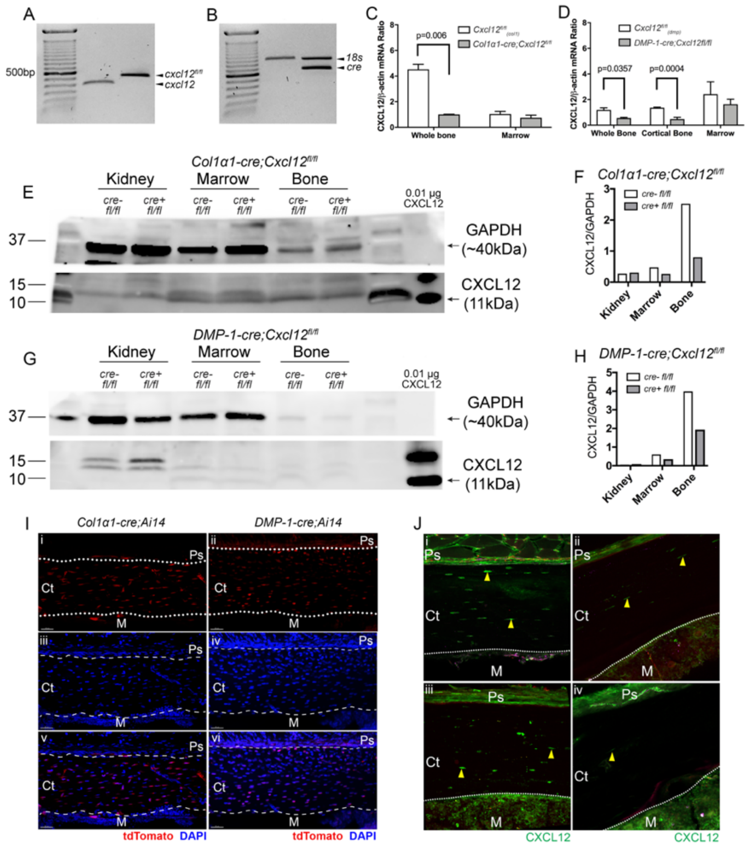

**Fig. S6.** Confirmation of *cxcl12* gene deletion driven by the Col1α1 and DMP-1 promoters. (A) DNA gel electrophoresis confirmed the presence of the floxed *cxcl12* gene (500bp) and (B) the *cre* transgene (750 bp) in 16-week-old mice. (C) *Col1α1-cre;Cxcl12*$^{fl/fl}$ mice exhibited a 75% decrease in whole bone CXCL12 mRNA expression. (D) *DMP-1-cre;Cxcl12*$^{fl/fl}$ mice exhibited a 60% reduction in whole bone CXCL12 mRNA expression and a reduction of 75% in cortical bone CXCL12 mRNA expression by qRT-PCR. (E, F) CXCL12 protein expression in whole bone was reduced by approximately 70% *Col1α1-cre;Cxcl12*$^{fl/fl}$ mice and (G, H) 50% in *DMP-1-cre;Cxcl12*$^{fl/fl}$ mice compared with their respective controls by Western blot. (I) Representative images of DAPI and tdTomato fluorescence in *Col1α1-cre;Ai14* and *DMP-1-cre;Ai14* mice. (i, ii) tdTomato, (iii, iv) DAPI, (v, vi) overlay. (J) Immunohistochemical detection of CXCL12 in cortical bone from (i) *Cxcl12*$^{fl/fl}_{(col1)}$, (ii) *Col1α1-cre;Cxcl12*$^{fl/fl}$, (iii) *Cxcl12*$^{fl/fl}_{(dmp)}$ and (iv) *DMP-1-cre;Cxcl12*$^{fl/fl}$ mice. CXCL12-expressing cells (green) in cortical bone in (i) *Cxcl12*$^{fl/fl}_{(col1)}$ and (iii) *Cxcl12*$^{fl/fl}_{(dmp)}$ control mice (yellow arrows). Fewer positively-stained osteocytes are observed in (ii) *Col1α1-cre;Cxcl12*$^{fl/fl}$ and (iv) *DMP-1-cre;Cxcl12*$^{fl/fl}$ mice (yellow arrows). Ps, periosteum; Ct, cortical bone; M, marrow. Scale bar = 30 µm.

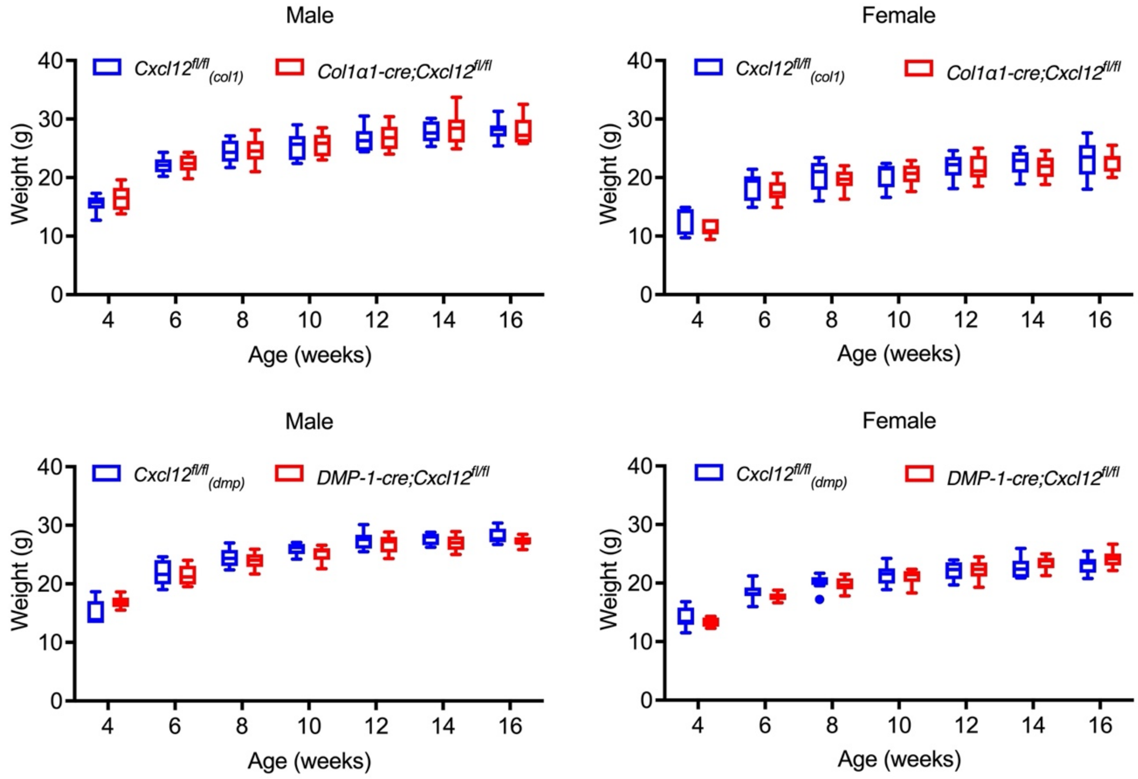

**Fig. S7.** Body weight of *Col1α1-cre;Cxcl12*$^{fl/fl}$ and *DMP-1-cre;Cxcl12*$^{fl/fl}$ mice during development. Body weight of male and female *Col1α1-cre;Cxcl12*$^{fl/fl}$ and *DMP-1-cre;Cxcl12*$^{fl/fl}$ mice, and their respective littermate controls *Cxcl12*$^{fl/fl}_{(col1)}$ and *Cxcl12*$^{fl/fl}_{(dmp)}$, from 4 to 16 weeks of age. N ≥ 7.

### SI Tables

**Table S1.** Geometric and biomechanical properties of the femur in female *Col1α1-cre;Cxcl12*$^{fl/fl}$ and *DMP-1-cre;Cxcl12*$^{fl/fl}$ and their respective controls.

| | ***Cxcl12***$^{fl/fl}_{(col1)}$ | ***Col1α1-cre;Cxcl12***$^{fl/fl}$ | ***Cxcl12***$^{fl/fl}_{(dmp)}$ | ***DMP-1-cre;Cxcl12***$^{fl/fl}$ |
|---|---|---|---|---|
| Length (mm) | 17.53±0.20 | 17.41±0.22 | ***15.94±0.03*** | ***16.11±0.09***[a] |
| Cortical Area (mm$^2$) | 0.846±0.037 | 0.780±0.069 | 0.856±0.036 | 0.849±0.022 |
| Cortical Thickness (mm) | 0.209±0.005 | 0.199±0.012 | 0.211±0.006 | 0.207±0.008 |
| Imin (mm$^4$) | 0.118±0.011 | 0.107±0.017 | 0.115±0.012 | 0.115±0.008 |
| Imax (mm$^4$) | 0.196±0.021 | 0.170±0.022 | 0.206±0.023 | 0.213±0.016 |
| J (mm$^4$) | 0.314±0.032 | 0.278±0.038 | 0.321±0.033 | 0.330±0.016 |
| Stiffness (N/mm) | 82.03±4.38 | 76.69±14.77 | 74.53±14.84 | 73.29±14.56 |
| Yield Load (N) | 11.70±0.88 | 10.81±1.33 | 12.52±1.50 | 11.30±0.67 |
| Peak Load (N) | 15.36±1.21 | 14.41±1.58 | 14.55±0.99 | 13.63±1.14 |
| Failure Load (N) | 15.15±1.27 | 13.23±2.450 | 12.83±1.389 | 12.51±1.47 |
| Displacement at Failure (mm) | 4.613±0.130 | 4.755±0.185 | 4.658±0.171 | 4.697±0.152 |
| P-Y Displacement (mm) | 0.372±0.127 | 0.544±0.206 | 0.490±0.146 | 0.593±0.106 |
| P-Y Work-to-Failure (N*mm) | 5.302±1.692 | 7.132±2.637 | 2.926±1.688 | 2.760±2.188 |
| Elastic Modulus (GPa) | 4.957±0.303 | 5.096±0.254 | 4.675±0.923 | 4.476±0.832 |
| Yield Stress (GPa) | 0.110±0.010 | 0.112±0.014 | ***0.120±0.010*** | ***0.106±0.007***[a] |
| Ultimate Stress (GPa) | 0.142±0.013 | 0.136±0.023 | 0.124±0.015 | 0.117±0.014 |

*Cxcl12*$^{fl/fl}_{(col1)}$ and *Cxcl12*$^{fl/fl}_{(dmp)}$ are Cre-negative littermate controls for the *Col1α1-cre;Cxcl12*$^{fl/fl}$ and *DMP-1-cre;Cxcl12*$^{fl/fl}$ mouse lines, respectively.
Imax, maximum area moment of inertia; Imin, minimum area moment of inertia; J, polar moment of inertia; P-Y, post-yield; N, newton; GPa, gigapascal
n=5-6 samples/group
[a]$p<0.05$ vs. respective control within sex by a two-tailed Student's t-test

**Table S2.** Geometric and biomechanical properties of the femur in male *Col1α1-cre;Cxcl12*$^{fl/fl}$ and *DMP-1-cre;Cxcl12*$^{fl/fl}$ and their respective controls.

| | ***Cxcl12***$^{fl/fl}_{(col1)}$ | ***Col1α1-cre;Cxcl12***$^{fl/fl}$ | ***Cxcl12***$^{fl/fl}_{(dmp)}$ | ***DMP-1-cre;Cxcl12***$^{fl/fl}$ |
|---|---|---|---|---|
| Length (mm) | 17.538±0.201 | 17.4067±0.224 | 15.992±0.039 | 15.958±0.064 |
| Cortical Area (mm$^2$) | 0.808±0.052 | 0.830±0.067 | 0.862±0.059 | 0.846±0.040 |
| Cortical Thickness (mm) | 0.196±0.008 | 0.198±0.0.009 | 0.198±0.010 | 0.196±0.009 |
| Imin (mm$^4$) | 0.111±0.014 | 0.122±0.018 | 0.128±0.017 | 0.128±0.013 |
| Imax (mm$^4$) | 0.230±0.033 | 0.220±0.026 | 0.245±0.027 | 0.247±0.021 |
| J (mm$^4$) | 0.340±0.045 | 0.342±0.044 | 0.374±0.044 | 0.374±0.031 |
| Stiffness (N/mm) | 68.58±14.97 | 67.17±15.0 | 78.75±12.25 | 72.26±7.75 |
| Yield Load (N) | 10.72±1.44 | 10.48±2.18 | 10.22±1.55 | 11.23±2.05 |
| Peak Load (N) | 14.24±1.77 | 15.8±2.12 | 15.08±2.19 | 14.38±0.72 |
| Failure Load (N) | ***13.49±0.96*** | ***15.34±1.35***[a] | 13.71±2.97 | 12.60±1.13 |
| Displacement at Failure (mm) | 4.408±0.134 | 4.440±0.271 | 4.263±0.138 | 4.220±0.164 |
| P-Y Displacement (mm) | 0.324±0.118 | 0.361±0.261 | 0.143±0.010 | 0.172±0.027 |
| P-Y Work-to-Failure (N*mm) | 4.455±2.081 | 5.212±4.007 | 3.793±2.188 | 2.131±1.702 |
| Elastic Modulus (GPa) | 4.419±0.761 | 3.931±0.525 | 4.371±0.588 | 3.897±0.538 |
| Yield Stress (GPa) | 0.105±0.010 | 0.095±0.018 | 0.120±0.016 | 0.103±0.018 |
| Ultimate Stress (GPa) | 0.132±0.009 | 0.139±0.012 | 0.121±0.019 | 0.116±0.018 |

*Cxcl12*$^{fl/fl}_{(col1)}$ and *Cxcl12*$^{fl/fl}_{(dmp)}$ are Cre-negative littermate controls for the *Col1α1-cre;Cxcl12*$^{fl/fl}$ and *DMP-1-cre;Cxcl12*$^{fl/fl}$ mouse lines, respectively.
Imax, maximum area moment of inertia; Imin, minimum area moment of inertia; J, polar moment of inertia; P-Y, post-yield; N, newton; GPa, gigapascal
n=5-6 samples/group
[a]$p<0.05$ vs. respective control within sex by a two-tailed Student's t-test

**Table S3.** Primers, Taqman probes and antibodies

| Category | Target | Description and Details | Manufacturer | Catalog # |
|---|---|---|---|---|
| **Primers** | *cxcl12* | Forward-5'- CTA CAC CTC CTC TAG GTA AAC CAG TCA GCC-3' | Invitrogen | A15612 |
| | | Reverse-5' GGA CAC CAG AAC CTT GAA ACT GACA-3' | Invitrogen | A15612 |
| | *cre* | Forward- 5'-CCT GGA AAA TGC TTC TGT CCT TTG CC-3' | Invitrogen | A15610 |
| | | Reverse-5'-GAG TTG ATA GCT GGC TGG TGG CAG ATG-3' | Invitrogen | A15610 |
| | *18S* | Forward- 5'-CAA GGA AGG CAG CAG GCG CGC AAA T-3' | Invitrogen | A15610 |
| | | Reverse-5'-TGC ACC ACC ACC CAC GGA ATC GAG AA-3' | Invitrogen | A15610 |
| **Taqman Probes** | CXCL12 | Assay #Mm00445553_m1 | Thermo Fisher | 4331182 |
| | 18S | Assay #Mm03928990_g1 | Thermo Fisher | 4331182 |
| | CXCR4 | Assay #Mm01996749_s1 | Thermo Fisher | 4331182 |
| | Runx2 | Assay #Mm00501584_m1 | Thermo Fisher | 4331182 |
| | Sp7 (Osx) | Assay #Mm04209856_m1 | Thermo Fisher | 4331182 |
| | Axin2 | Assay #Mm00443610_m1 | Thermo Fisher | 4331182 |
| **Antibodies** | | *Details* | | |
| WB | CXCL12 | 1:1000 anti-mouse | Cell Signal | 5558 |
| | GAPDH | 1:2500 anti-mouse | Cell Signal | 4691 |
| | GSK3β | 1:1000 anti-mouse | Cell Signal | 4058 |
| | pGSK3β | 1:1000 anti-mouse | Cell Signal | 5558 |
| | Akt | 1:1000 anti-mouse | Cell Signal | 4691 |
| | pAkt | 1:1000 anti-mouse | Cell Signal | 4058 |
| | β-catenin | 1:1000 anti-mouse | Cell Signal | 9582 |
| | pβ -catenin | 1:1000 anti-mouse | Cell Signal | 9561 |
| | β-actin | 1:1000 anti-mouse | Cell Signal | 4970 |
| IHC | CXCL12 | 1:100 rabbit anti-mouse | Santa Cruz | 28876 |
| | Osteocalcin | 1:100 goat anti-mouse | Fisher Sci | PA1-85754 |
| | Cathepsin K | 1:100 rabbit anti-mouse | Abcam | ab19027 |
| | Alexafluor-488 | 1: 300 donkey anti-rabbit | Fisher Sci | A-21206 |
| | Alexafluor-594 | 1:300 goat anti-rabbit | Jackson IR | 705-585-003 |
| FACS | Ter119-APC | clone TER-119 | eBioscience | Ref#17-5921-82 |
| | CD71-PECy7 | clone RI7217 | Biolegend | Ref#113812 |
| | CD45-PE | clone 30-F11 | eBioscience | Ref#12-0451-82 |
| | CD3-PE | clone 17A2 | Biolegend | Ref#100206 |
| | B220-PE | clone RA3-6B2 | Biolegend | Ref#103208 |
| | CD19-PE | clone 6D5 | Biolegend | Ref#115508 |
| | Gr-1-PE | clone RB6-8C5 | Biolegend | Ref#108408 |
| | Cd11b-PE | clone M1/70 | Biolegend | Ref#101208 |
| | 7-AAD | 1:50 | Thermo Fisher | 00-6993-50 |

**SI Movies**

**Movie SI.** Rotating three-dimensional image dataset of a tibia at mid-shaft from a normally ambulating CXCL12dsRed reporter mouse, which expresses DsRed-Express2 from the endogenous *cxcl12* mouse promoter and is counterstained with DAPI. The volume of interest consists of cortical bone with singly encased osteocytes distributed throughout the matrix, which is depicted in gray originating from the second harmonic generation (SHG) signal. Osteocytes appear blue with low expression of DsRed. Bone marrow cells of heterogeneous size and shape are densely packed in an irregular pattern next to the endocortical surface of the cortex; they appear blue with varying levels of expression of DsRed. DsRed is visible primarily in marrow cells.

**Movie S2.** Rotating three-dimensional image dataset of an exogenously-loaded tibia at mid-shaft from a CXCL12dsRed reporter mouse, which expresses DsRed-Express2 from the endogenous *cxcl12* mouse promoter and is counterstained with DAPI. The volume of interest consists of cortical bone with singly encased osteocytes distributed throughout the matrix, which is depicted in gray originating from the second harmonic generation (SHG) signal. Osteocytes appear blue with high expression of DsRed as a result of increased *in vivo* mechanical loading. Bone marrow cells of heterogeneous size and shape are densely packed in an irregular pattern next to the endocortical surface of the cortex; they appear blue with varying levels of expression of DsRed. The periosteum and adjacent muscle are situated at the periosteal surface of the cortex. Cells in the periosteum, presumably of osteogenic potential, appear red. DsRed is visible throughout the marrow, cortical bone and periosteum.
